# The SWI/SNF PBAF complex facilitates REST occupancy at repressive chromatin

**DOI:** 10.1101/2024.08.23.609212

**Authors:** Elena Grossi, Christie B. Nguyen, Saul Carcamo, Shannon Moran, Valentina Kirigin Callaú, Dan Filipescu, Dan Hasson, Emily Bernstein

## Abstract

Multimeric SWI/SNF chromatin remodelers assemble into discrete conformations with unique complex functionalities difficult to dissect. Distinct cancers harbor mutations in specific subunits, altering the chromatin landscape, such as the PBAF-specific component ARID2 in melanoma. Here, we performed comprehensive epigenomic profiling of SWI/SNF complexes and their associated chromatin states in melanoma and melanocytes and uncovered a subset of PBAF-exclusive regions that coexist with PRC2 and repressive chromatin. Time-resolved approaches revealed that PBAF regions are generally less sensitive to ATPase-mediated remodeling than BAF sites. Moreover, PBAF/PRC2-bound loci are enriched for REST, a transcription factor that represses neuronal genes. In turn, absence of ARID2 and consequent PBAF complex disruption hinders the ability of REST to bind and inactivate its targets, leading to upregulation of synaptic transcripts. Remarkably, this gene signature is conserved in melanoma patients with ARID2 mutations. In sum, we demonstrate a unique role for PBAF in generating accessibility for a silencing transcription factor at repressed chromatin, with important implications for disease.

## Introduction

The mammalian SWItch/Sucrose Non-Fermentable (SWI/SNF) complexes are multimeric chromatin remodelers that utilize the energy of ATP hydrolysis to mobilize nucleosomes and provide an accessible chromatin substrate for transcription factors (TFs) and RNA Polymerase II (Pol II) machinery^1,2^. Recent structural and biochemical studies have unveiled important details regarding SWI/SNF complex composition, as well as how these complexes engage the nucleosome octamer^3–6^. Nevertheless, the mechanisms by which chromatin remodelers that lack sequence-specific DNA binding ability recognize specific genomic targets, remains unclear. While *in vitro* studies point towards histone modifications and chromatin states dictating SWI/SNF binding and remodeling activity^7^, recent *in vivo* evidence suggests a more dynamic mechanism, whereby these complexes scan the genome, transiently engaging the nucleosome, and productively remodel chromatin only when interacting with specific TFs^8,9^. Importantly, various TFs rely on distinct families of chromatin remodeling complexes (*e.g.* SWI/SNF or ISWI) for proper genomic binding^10^, highlighting a co-dependency between these chromatin components.

Further complicating the functional dissection of individual SWI/SNF complexes is the modular nature of these macromolecular assemblies, that generally exist in three distinct conformations, known as canonical BAF (cBAF), non-canonical BAF (ncBAF), and Polybromo-associated BAF (PBAF)^6^. These three complex subfamilies are characterized by unique components, as well as several common core subunits including an ATPase module, whose catalytic activity is continuously required to maintain chromatin accessibility, especially at enhancer elements^11–14^. SWI/SNF remodeling, particularly by cBAF, also mediates the rapid eviction of Polycomb Repressive Complexes (PRC1/2) and in turn, their associated histone marks (H2AK119ub and H3K27me3, respectively), establishing accessible chromatin within minutes of its occupancy^15,16^. Moreover, different SWI/SNF complexes preferentially bind distinct regulatory regions of the genome, with PBAF mainly enriched at promoters and cBAF primarily occupying enhancers^17^. However, the molecular basis governing these preferences remains unclear, as most studies elucidating SWI/SNF mechanisms of action have focused on common SWI/SNF complex subunits (*e.g.,* the enzymatic subunit BRG1/SMARCA4), and are therefore unable to discriminate between distinct complex subfamilies or uncover potential functional differences.

Understanding how each SWI/SNF remodeling complex operates is of paramount importance, particularly in the context of human disease. While mutations in SWI/SNF subunits are collectively found in >20% of human cancers, genetic alterations in individual components are enriched in distinct tumors rather than randomly distributed across cancer types, suggesting lineage- and subunit-specific functions^18–20^. For instance, the PBAF-specific component ARID2 is considered a mutational ‘cancer driver’ in melanoma^21–24^, an aggressive skin cancer derived from melanocytes. ARID2 alterations are present in 18% of melanoma samples, according to the Cancer Genome Atlas (TCGA) database^25–27^, and are largely missense or nonsense with very few hot spots, pointing towards a tumor suppressive role. Interestingly, ARID2 mutations have been detected in early melanoma lesions^22^, yet are also enriched in melanoma metastases^28,29^, suggesting that this factor might be important at different stages of melanoma onset and progression. Recent evidence from our group showed that ARID2 loss-of-function in melanoma impairs PBAF complex assembly, in line with the structural role of ARID subunits in complex formation^6^. This leads to BAF genomic redistribution, ultimately resulting in activation of transcriptional programs associated with melanoma cell dissemination^30^. Nevertheless, it remains unclear whether PBAF utilizes unique molecular mechanisms distinct from other SWI/SNF complexes that are essential in specific developmental and/or tumoral contexts.

To address this question, we carried out comprehensive epigenomic profiling of the SWI/SNF-associated chromatin landscape in melanoma cells and primary normal human epidermal melanocytes (NHEM). We found that PBAF-enriched regions exhibit lower levels of chromatin accessibility than cBAF and ncBAF sites, and a subset of PBAF occupied regions are unexpectedly enriched for H3K27me3 and the PRC2 complex. By employing a SWI/SNF chemical inhibitor in a time-resolved fashion, we further revealed that PBAF regions are less sensitive to ATPase-dependent remodeling than those of BAF, with different response rates associated with distinct sets of TFs. Among them, we identified the transcriptional repressor REST (<u>RE</u>1 <u>S</u>ilencing <u>T</u>ranscription Factor) as a PBAF-specific TF enriched at inactive regions, and whose chromatin occupancy is dependent on PBAF. Accordingly, REST target genes are derepressed upon ARID2 depletion, as well as in human melanoma tumors bearing ARID2 mutations, unveiling a molecular crosstalk between a chromatin remodeler with modest remodeling activity and a repressive TF at inactive chromatin.

## Results

### A subset of PBAF-enriched regions associates with repressed chromatin

We previously demonstrated that in the absence of functional PBAF complexes in melanoma, BAF redistributes genome-wide, ultimately affecting chromatin accessibility and TF binding^30^. This suggests a delicate balance between SWI/SNF subcomplexes and their associated TFs. To characterize genomic regions uniquely regulated by SWI/SNF complexes, as well as their associated epigenetic landscapes, we performed genomic mapping studies in melanoma cells and NHEM (**Fig. 1A, B**). PBAF-enriched genomic regions were defined as loci bound by the PBAF-exclusive subunit ARID2, but lacking SS18, a SWI/SNF subunit present only in cBAF and ncBAF complexes^6^. Shared regions were characterized by occupancy of both ARID2 and SS18, while cBAF/ncBAF (referred to as BAF) showed exclusive enrichment for SS18 (**Fig. 1A, B; Table S1**).

**Figure 1.**
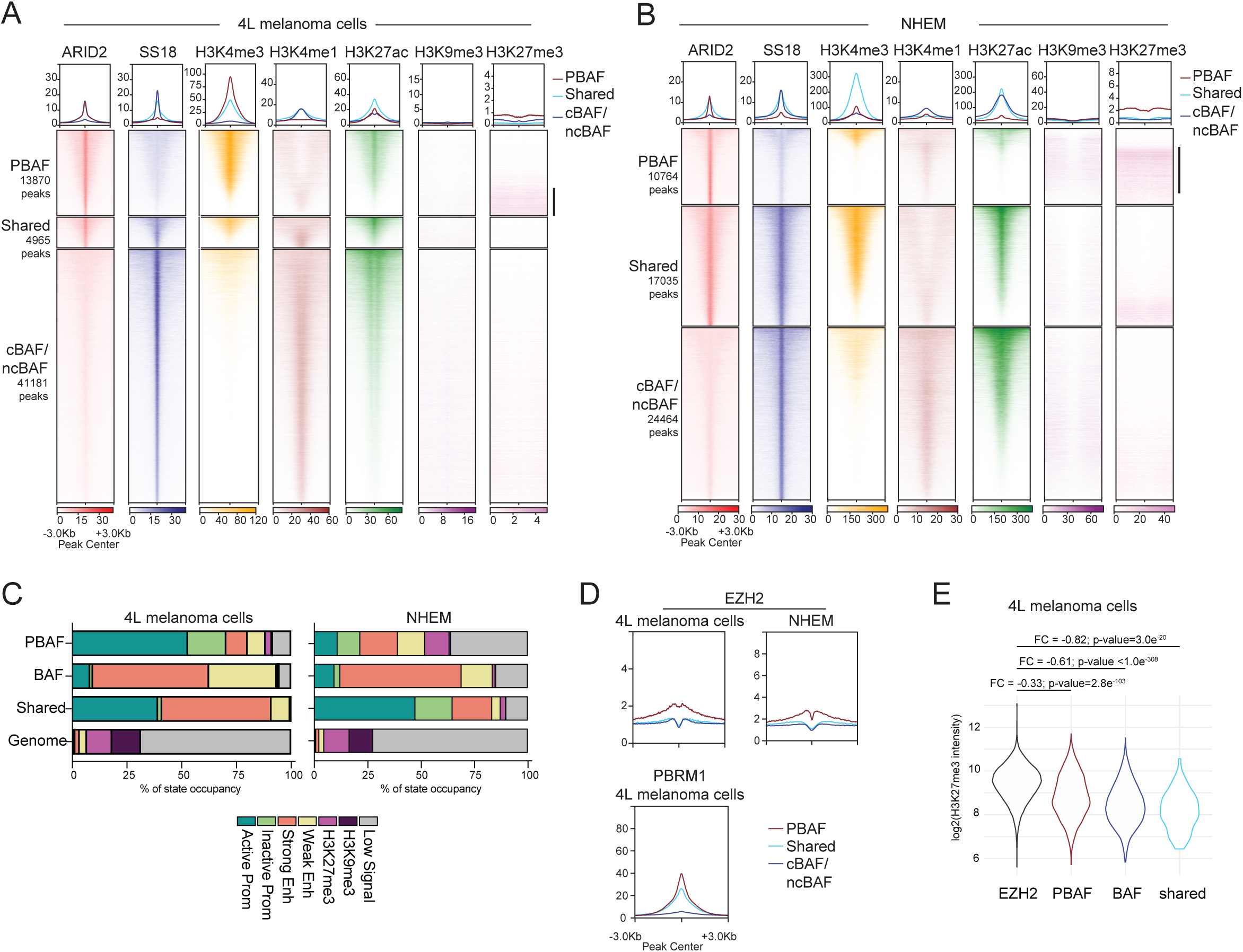
A subset of PBAF-enriched regions associates with repressed chromatin. (A-B) Heatmaps displaying the genomic distribution of the SWI/SNF components ARID2 and SS18, as well as of multiple histone marks, at PBAF-, Shared- or BAF-(cBAF/ncBAF-) enriched regions sorted by H3K27ac signal in (A) 4L melanoma cells or (B) NHEM (see Methods). Black sidebars highlight H3K27me3-rich PBAF regions. (C) Distribution of chromatin states associated with different SWI/SNF subcomplexes and compared to their distribution in the genome of 4L melanoma cells and NHEMs. (D) Metagene profiles of EZH2 and PBRM1 in 4L melanoma cells and EZH2 in NHEM at PBAF-, Shared, or BAF-enriched regions as defined in (A). (E) Violin plot showing the distribution of H3K27me3 signal in 4L melanoma cells at EZH2-(n=37168), PBAF-(n=1241), BAF-(n=1707) enriched or shared (n=58) regions. Only regions overlapping H3K27me3 peaks were selected (see Fig. S1F for unedited data). A pseudocount was added for logarithmic scale visualization. Mean signal for each group was used to calculate fold change (FC) values. Significance was calculated by Mann-Whitney U test.

By profiling a variety of active and repressive histone modifications in two melanoma cell lines with distinct driver mutations, 113/6-4L (herein referred to as 4L; BRAF^V600E^) and SKmel147 (NRAS^Q61R^), we confirmed the observation that PBAF is enriched at active promoters (marked by H3K4me3) but less so at functional enhancers (marked by H3K4me1), while BAF primarily binds enhancers, and shared regions encompass both regulatory elements (**Fig. 1A, S1A, B**)^30^. Interestingly, the PBAF binding profile was somewhat distinct in NHEM, displaying enrichment at both proximal and distal regulatory regions of the genome (**Fig. S1B**). In addition, H3K4me3 levels at PBAF-exclusive regions appeared drastically reduced in NHEM, whereas they remained high at shared sites. While the reduction in H3K4me3 might be associated with the global decrease of this histone modification upon cellular differentiation^31,32^, the concomitant increased proportion of shared regions in NHEM *vs.* melanoma might also indicate that in differentiated cells, the PBAF complex relies on the presence of BAF to exert its role at promoters (**Fig. 1B, S1B**). Nonetheless, in both normal and tumor cells, a large fraction of these regulatory elements was marked by H3K27ac, with shared regions (i.e., BAF/PBAF-bound promoters) exhibiting the highest enrichment.

Notably, at a subset of PBAF-enriched regions lacking H3K27ac, we unexpectedly identified enrichment of H3K27me3, a mark deposited by PRC2 (**Fig. 1A, B, S1A**). This feature was not exclusive to tumor cells, as we observed even higher H3K27me3 enrichment at PBAF-exclusive regions in NHEM compared to melanoma cells (**Fig. 1B**). This was not the case for H3K9me3, a histone modification associated with constitutive heterochromatin (**Fig. 1A, B**, **S1A**). Consistently, chromatin state models generated with ChromHMM^33^ revealed that ∼3% of PBAF peaks in melanoma cells (4L and SKmel147), and 11% in melanocytes, occupy regions exclusively marked by H3K27me3 (**Fig. 1C, S1C, D**). In comparison, only 0.2-0.8% and 1% of BAF sites (in melanoma and melanocytes, respectively) colocalize with this repressive mark. The higher enrichment found in NHEM might reflect the more repressed epigenome of differentiated cells^34^, in particular at regulatory regions of the genome, as some H3K27me3 was also observed in shared and BAF-enriched regions in NHEM (**Fig. 1B, C**). These results, obtained by H3K27me3 CUT&RUN, were further validated by ChIP-seq using an unrelated H3K27me3 antibody (**Fig. S1E, F**). Taken together, these data suggest that PBAF and PRC2 may co-exist at certain chromatin loci, in contrast with the previously described antagonistic relationship between the BAF and PRC complexes^15,16^.

Next, we aimed to confirm occupancy of the PRC2 complex at these H3K27me3-marked regions. To do so, we mapped the genomic binding of the PRC2 catalytic subunit EZH2 and found it was indeed enriched at PBAF-specific regions in both normal and tumor cells (**Fig. 1D, S1E, G**). As expected, PBAF is associated with far fewer H3K27me3 sites than EZH2, yet a subset of PBAF regions present similar levels of H3K27me3, while BAF and shared regions display greatly reduced levels (**Fig. 1E, S1H**). Lastly, ChIP-seq for the core catalytic SWI/SNF component BRG1 and the PBAF-specific subunit PBRM1 confirmed that PBAF-enriched repressed regions are indeed bound by fully assembled PBAF complexes (**Fig. 1D**, **S1A, E, G**), as PBRM1 is one of the last subunits to be assembled into PBAF and no partially assembled complexes containing PBRM1 have been identified^6^. These data collectively indicate that PBAF can co-localize with PRC2 at a subset of repressed genomic regions devoid of the BAF complex.

### PBAF-enriched regions exhibit lower levels of chromatin accessibility than those of BAF

Since PRC eviction from chromatin can be mediated by the ATPase-dependent remodeling activity of the SWI/SNF complex^15^, we tested chromatin accessibility levels across the SWI/SNF regions described above in both normal and transformed cells. Interestingly, we found that accessibility at PBAF-enriched chromatin regions is generally lower compared to shared or BAF sites in all melanoma cell lines examined, as well as in NHEM (**Fig. 2A, B, S1A)**. We then asked whether occupancy levels of either BAF or PBAF complexes correlated with accessibility status at their respective bound regions. We found that while SS18 binding presents a strong, positive correlation with chromatin accessibility at BAF loci (R= 0.64), correlation values are much lower for ARID2 binding (R=0.31), indicating that PBAF presence alone is not sufficient to efficiently open chromatin (**Fig. 2C**).

**Figure 2.**
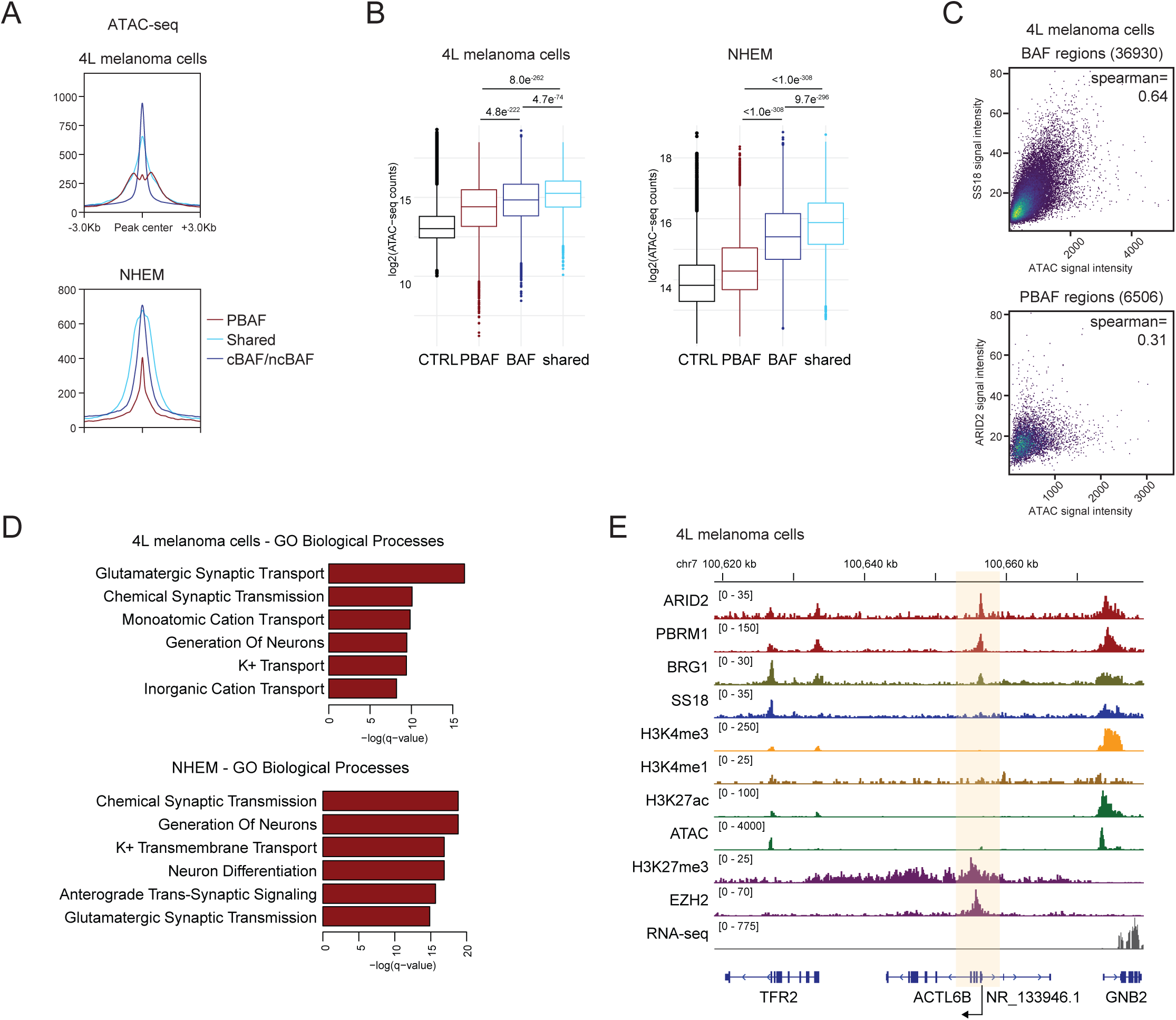
PBAF-enriched regions present a lower level of remodeling activity. (A) Metagene plots displaying ATAC-seq levels at PBAF-, Shared, or BAF-enriched regions in 4L melanoma cells and NHEM (B) Box plot representing chromatin accessibility levels in 4L melanoma cells (left panel) or NHEMs (right panel) at PBAF-, Shared or BAF-enriched regions. Control (CTRL) sites include ATAC peaks not overlapping with any SWI/SNF region. Significance was calculated by Mann-Whitney U test. (C) Scatter plot showing correlation levels between SS18 or ARID2 levels and ATAC signal at their respective regions. To ensure that only fully assembled SWI/SNF complexes were represented in this analysis, only SS18 and BRG1 or ARID2, PBRM1 and BRG1 overlapping peaks were included as BAF and PBAF regions, respectively. (D) Biological Processes GO analysis of genes localized at repressed PBAF chromatin regions (cluster #1 of Fig. S2A, B) in melanoma cells or NHEMs. (E) Genome browser snapshot showing occupancy of SWI/SNF and PRC2 components, as well as histone marks at *ACTL6B* genomic locus in 4L melanoma cells.

While low ATAC-seq signal appears to be a common feature of PBAF-enriched regions, only a subset of these sites displays enrichment of EZH2 and H3K27me3, as shown by k-means clustering of PBAF-enriched peaks in melanoma cells and NHEM (**Fig. S2A, B**). To understand the biological relevance of these PBAF-PRC2 regions, we associated such sites (811 peaks in 4L and 2263 in NHEM) with the nearest gene. RNA-seq analysis in normal and tumor cells revealed that these genes display very low expression levels, in line with their associated chromatin state (**Fig. S2C, D**). Moreover, Gene Ontology (GO) analysis identified significant enrichment for terms related to synaptic transmission, ion channel function, and neuron differentiation in both normal and tumor cells (**Fig. 2D** and **Table S2**). Among these PBAF-occupied, ATAC-low, H3K27me3-enriched repressed genes associated with neuronal function is *ACTL6B*, a member of the neuronal BAF (nBAF) complex, a lineage-specific BAF assembly that is necessary for neurogenesis^35^ (**Fig. 2E**). Interestingly, melanocytes share a common neural crest precursor with neuronal cells and silencing of neuronal-related genes is a critical step in the establishment of the melanocyte lineage, while their deregulation has been associated with increased malignancy^36–39^. Taken together, these results indicate that PBAF binding at its genomic targets, in the absence of BAF co-occupancy, results in a lower level of chromatin remodeling inferred by chromatin accessibility, thus possibly preserving an inactive or poised status of key developmental genes.

### SWI/SNF complex subfamilies present distinct remodeling dynamics

To better understand the differences in remodeling activity of distinct SWI/SNF complexes, we utilized a selective BRG1/BRM ATPase inhibitor (BRM014), which has recently allowed the study of SWI/SNF activity at unprecedented resolution^12–14,40^. We treated 4L melanoma cells with 1 μM of BRM014 for 10 minutes, 1 hour, or 6 hours, to temporally resolve the remodeling activity of different SWI/SNF complexes in a short timeframe using ATAC-seq as readout. Importantly, this inhibitor does not affect SWI/SNF chromatin occupancy, nor impacts melanoma proliferation at these time points (**Fig. S3A, B**).

Unbiased analysis and k-means clustering of altered ATAC peaks revealed that upon ATPase inhibition, chromatin accessibility at distinct genomic regions is affected at different rates (**Fig. S3C**). For example, a large fraction of ATAC peaks is immediately affected by inhibitor treatment, quickly reducing accessibility levels within the first 10 minutes (clusters #2, #4); other clusters lose chromatin opening at a slower rate (clusters #3, #5), while a minimal set of regions increases accessibility (cluster #1). Interestingly, these patterns mirror what was reported in murine models^12^, suggesting conserved mechanisms in mammals.

By integrating altered ATAC-seq peaks across all time points, we observed a large fraction of regions with reduced accessibility (37%), while gained peaks appear to be a minority (0.7%), and the majority represented by constant regions, which may require other chromatin remodeling factors or prolonged inhibition (**Fig. 3A**). However, when we overlayed these regions with SWI/SNF occupancy, we observed that while SS18 and BRG1 are preferentially bound to decreased ATAC peaks, ARID2 is not. In fact, when we assigned ATAC-seq peaks to either BAF, shared, or PBAF regions, we observed that while BAF and shared sites largely overlap with regions that decrease in accessibility (64.5% and 42.3%, respectively), accessibility at PBAF-enriched loci was reduced in only 6.2% of cases (**Fig. 3B**). This suggests that genomic regions exclusively bound by PBAF are less sensitive to ATP-dependent remodeling inhibition.

**Figure 3.**
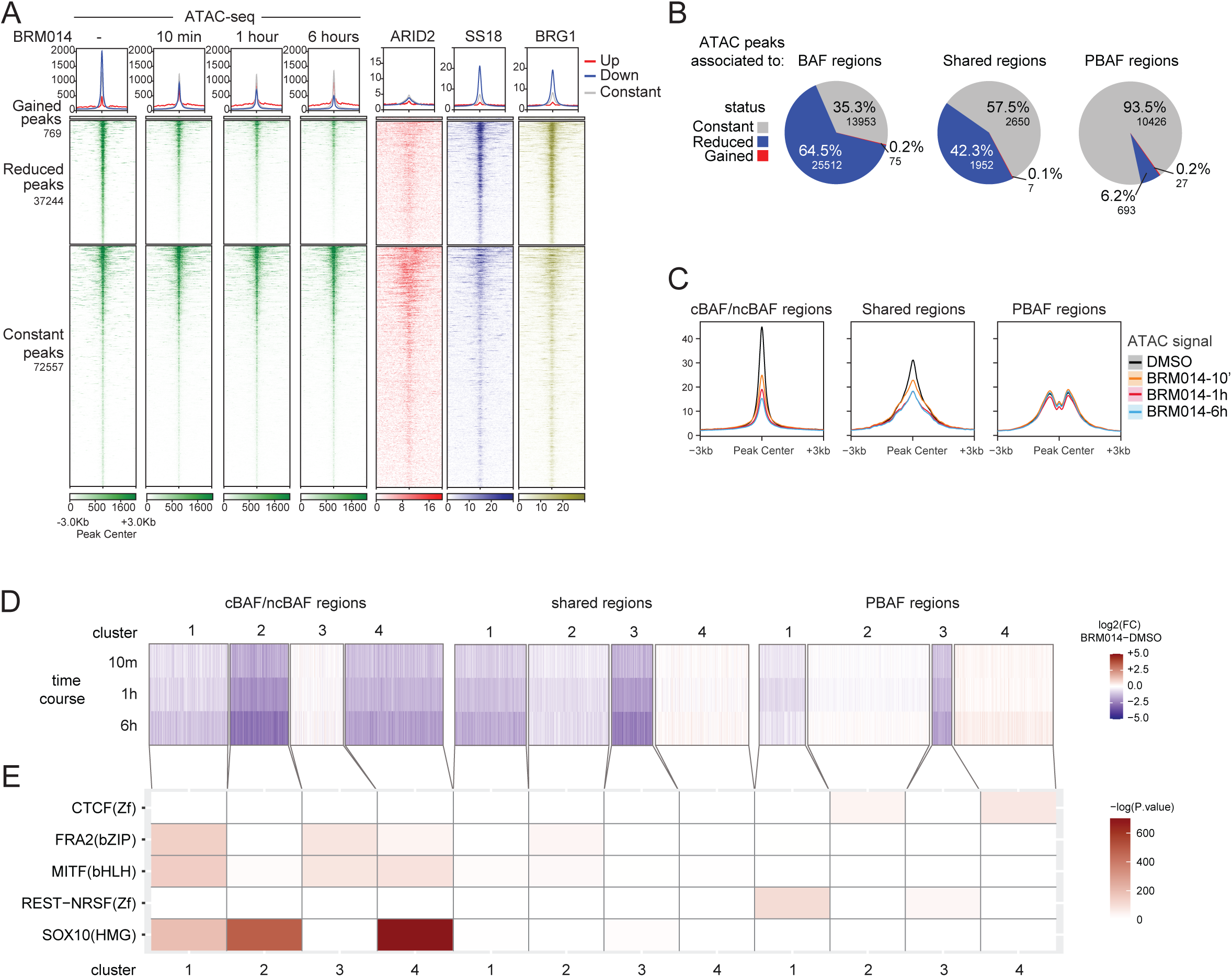
Different SWI/SNF subcomplexes display distinct remodeling dynamics. (A) Heatmap representing accessibility signal in 4L melanoma cells at differentially occupied ATAC regions upon BRM014 treatment, as determined by DiffBind analysis (see Methods). ChIP-seq signal for ARID2, SS18 and BRG1 components is also shown. (B) Pie chart illustrating the percentage of BAF, shared or PBAF regions associated to constant, reduced or gained ATAC peaks, as defined in (A). (C) Metagene profiles showing accessibility changes in 4L melanoma cells at BAF, shared or PBAF regions at different time points of BRM014 treatment. (D) Heatmap illustrating changes in accessibility signal in 4L melanoma cells at BAF, shared or PBAF regions during BRM014 treatment. K-means clustering was used to unbiasedly group regions in four clusters. Note that all ATAC regions were included in this analysis, regardless of the significance of the change (E) Heatmap displaying enrichment of selected TF binding motifs from publicly available datasets at different clusters of ATAC regions, as defined in (D). See complete list in Table S3.

Next, we examined the dynamics of remodeling across the different time points and the rate of accessibility changes for each SWI/SNF complex. At BAF-enriched regions, ATAC-seq signal is dramatically affected after only 10 minutes of BRM014 treatment and continues decreasing over time, likely reflecting the dependency on ATPase-mediated remodeling of enhancer elements (**Fig. 3C**). Shared BAF/PBAF co-occupied regions lose accessibility at early and intermediate time points but then stabilize at longer stages, possibly reflecting compensation mechanisms by other chromatin remodelers (*e.g.* EP400/TIP60) at promoter regions^14^. In contrast, PBAF-only regions present minimal to no reduction in ATAC signal upon BRM014 treatment, and this minor loss of accessibility occurs with a slower kinetics, requiring 1 hour of treatment.

We next asked whether co-occupancy of distinct TFs may play a role in response to ATPase inhibition, as they have been associated with specific sensitivities to ATPase-mediated remodeling. For instance, binding of the pluripotency TF OCT4 is rapidly reduced upon SWI/SNF catalytic inhibition in mouse embryonic stem cells (mESC), while CTCF is unaffected^12^. We therefore clustered SWI/SNF-associated ATAC peaks (both constant and affected regions) according to their respective complex and the rate of accessibility change at different time points and performed TF motif enrichment analysis, identifying differences within and between different complexes (**Fig. 3D, E** and **Table S3**). For example, SOX10 is exclusively found at ATPase-dependent, BAF-specific regions, while the MITF and FOSL2 (AP1) motifs are present at both shared and BAF regions but mostly enriched at loci unaffected or mildly altered upon BRM014 treatment. Intriguingly, PBAF regions displayed a strikingly different set of enriched TF motifs, including the insulator protein CTCF, characterizing the regions that gain accessibility upon BRM014 treatment, and the transcriptional repressor REST, which demarcates the small portion of ATPase-dependent PBAF regions (**Fig. 3D, E**).

To understand whether distinct TFs can affect the remodeling activity of different SWI/SNF complexes, we analyzed data from a recently published study performed in murine mammary epithelial cells^13^, in which PBAF-associated elements were found to be even less sensitive to BRM014 treatment compared to our study (0.1% after a 6-hour treatment). Motif enrichment analysis revealed that BAF-bound sites were associated with the AP1 family of TFs, in line with the role of this TF at enhancers and consistent with our previous results in melanoma cells^30^ (**Fig. S3D-E**). In contrast, PBAF-specific regions were enriched for the TF RONIN/GFY, while no REST motif was found at these sites, possibly explaining the lack of BRM014-sensitive PBAF regions in this cell system. Together, these results suggest that TF composition might influence the remodeling susceptibility of different genomic regions and constitute a discriminating feature to predict how chromatin sites respond to SWI/SNF remodeling inhibition.

### The transcription factor REST is preferentially associated with the PBAF complex

Our data point towards different remodeling sensitivities linked to distinct sets of TFs. We previously reported PBAF preference for the REST motif^30^. To further investigate the association between REST and SWI/SNF complexes, we performed ChIP-seq for this TF in melanoma cells and NHEM and confirmed its enrichment at PBAF-exclusive regions (**Fig. 4A**). Given that REST binds a relatively low number of regions (1556 peaks in melanocytes and 3081 in 4L melanoma cells), the subset of chromatin sites co-occupied by REST and PBAF is small, but it corresponds to the most repressed regions bound by PBAF (**Fig. S4A**). CTCF ChIP-seq in NHEM showed that this TF is localized at another well-defined and non-overlapping subset of PBAF sites, with these two TFs combined covering >25% of PBAF-occupied chromatin regions.

**Figure 4.**
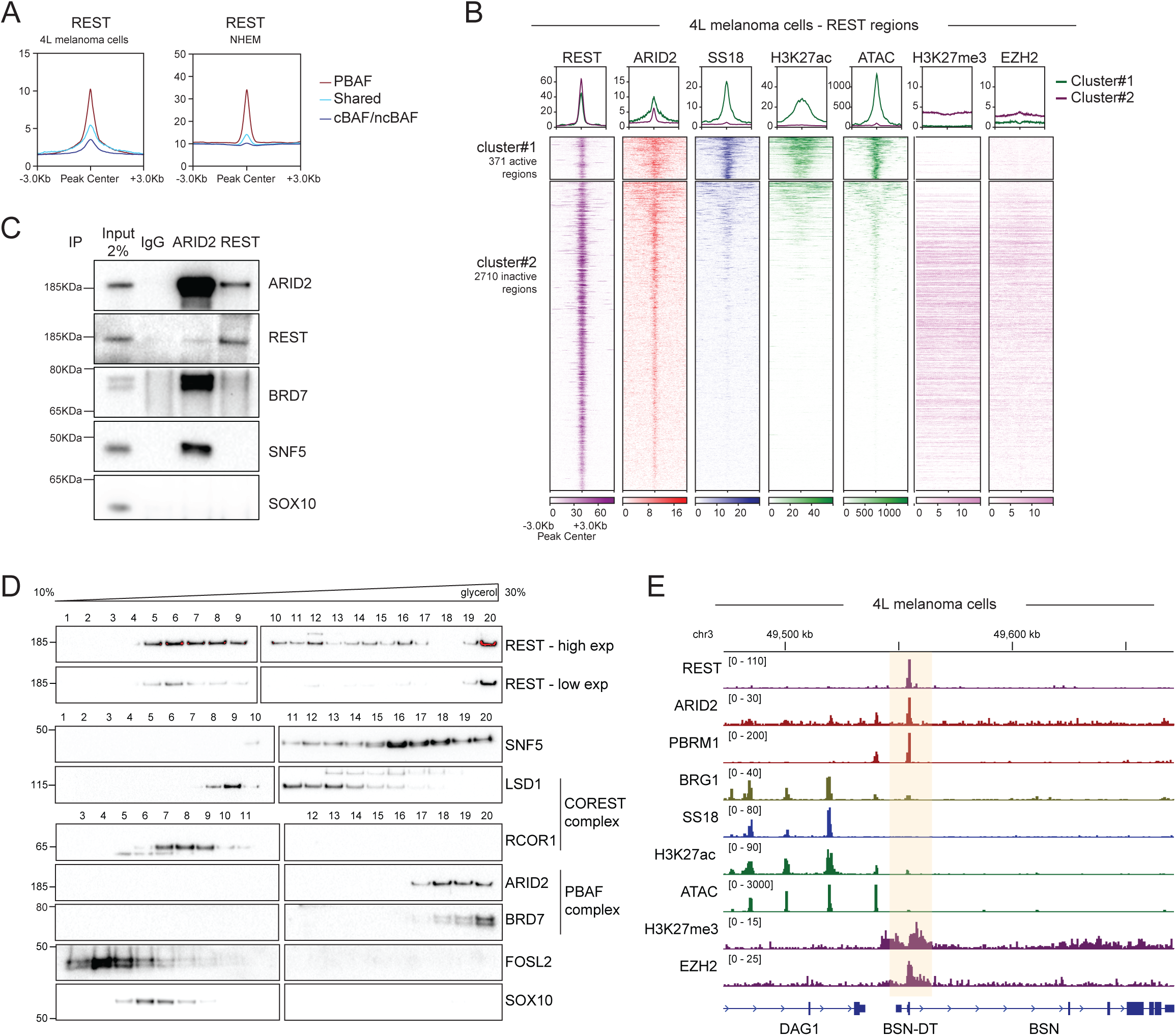
REST is preferentially associated with the PBAF complex. (A) Metagene profile of REST ChIP-seq in 4L cells and NHEMs at PBAF-, shared and BAF-enriched regions, as defined in Fig. 1A-B. (B) Heatmap displaying REST peaks grouped by k-means clustering, defining active (cluster#1) and inactive (cluster#2) regions in 4L melanoma cells. (C) Western blot analysis of ARID2 or REST immunoprecipitated chromatin extracts in 4L melanoma cells. (D) Immunoblot analysis of glycerol gradient sedimentation from SKmel147 melanoma cells. Different blots performed using the same protein extracts are shown. (E) Genomic browser snapshot illustrating occupancy of SWI/SNF and PRC2 components, histone marks and REST at *BSN* neuronal genes in 4L melanoma cells.

Interestingly, REST-bound regions in mESC were previously characterized by their unique response to SWI/SNF chemical inhibition, as they lose accessibility and TF binding at slower rates compared to regions bound by activating TFs^12^. We confirmed this slow dynamic in our human cell model (**Fig. S4B**) and linked this distinctive response to the presence of PBAF but no other SWI/SNF complexes (**Fig. 3E**). This connection is more evident when analyzing REST regions; only a limited number of REST peaks are associated with active chromatin marks and the BAF complex, as shown by SS18 binding, in line with its role as transcriptional repressor (**Fig. 4B**, **S4C**). In turn, the majority of REST sites overlap with ARID2 and repressed chromatin marks, such as H3K27me3 and EZH2, indicating that the PBAF complex and REST are functionally connected at inactive regions.

Next, we performed reciprocal co-immunoprecipitation (co-IP) using chromatin extracts to evaluate the physical association between PBAF and REST and confirmed that REST pull down enriches for ARID2 and *vice versa* (**Fig. 4C**). This interaction appears to be specific, as SOX10 is not enriched (**Fig. 4C**). Notably, by glycerol gradient sedimentation we observed that REST was enriched at both lighter and heavier fractions, whereas other TFs (such as SOX10 and FOSL2) occupy only the lighter fractions, suggesting that REST can form multiple stable macromolecular complexes (**Fig. 4D**). In line with our co-IP results, REST and ARID2 co-migrated to the heaviest fractions of the gradient, indicating that PBAF might stably interact with this repressive TF (**Fig. 4D**).

Taken together, these data point towards an association between the PBAF complex and REST at repressed PRC2-marked chromatin regions. REST targets related to synaptic function, such as *BSN* and *CHRNB2*, clearly exemplify these regions, presenting binding for PBAF but not BAF subunits, low enrichment of active marks and reduced accessibility, as well as PRC2 occupancy (**Fig. 4E, S4D)**.

### PBAF loss impairs REST binding at repressed regions

Given the above findings, we sought to investigate whether REST function is affected by loss of the PBAF complex. We therefore deleted ARID2 via CRISPR-Cas9 in melanoma cells and NHEM, as we and others showed that loss of ARID2 disrupts PBAF assembly^6,30^. Western blot analysis of chromatin extracts revealed that ARID2 knockout (KO) and the consequent PBAF complex disruption (demonstrated by loss of the PBAF-specific component PBRM1), reduced REST levels in chromatin without affecting other SWI/SNF subunits, such as SNF5/SMARCB1 (**Fig. 5A, B, S5A**). Interestingly, CTCF levels were not significantly affected in ARID2 KO cells (**Fig. 5A, B**). We therefore speculated that the weak remodeling activity of PBAF is in fact required to enable the binding of REST, but not CTCF, to its target chromatin sites.

**Figure 5.**
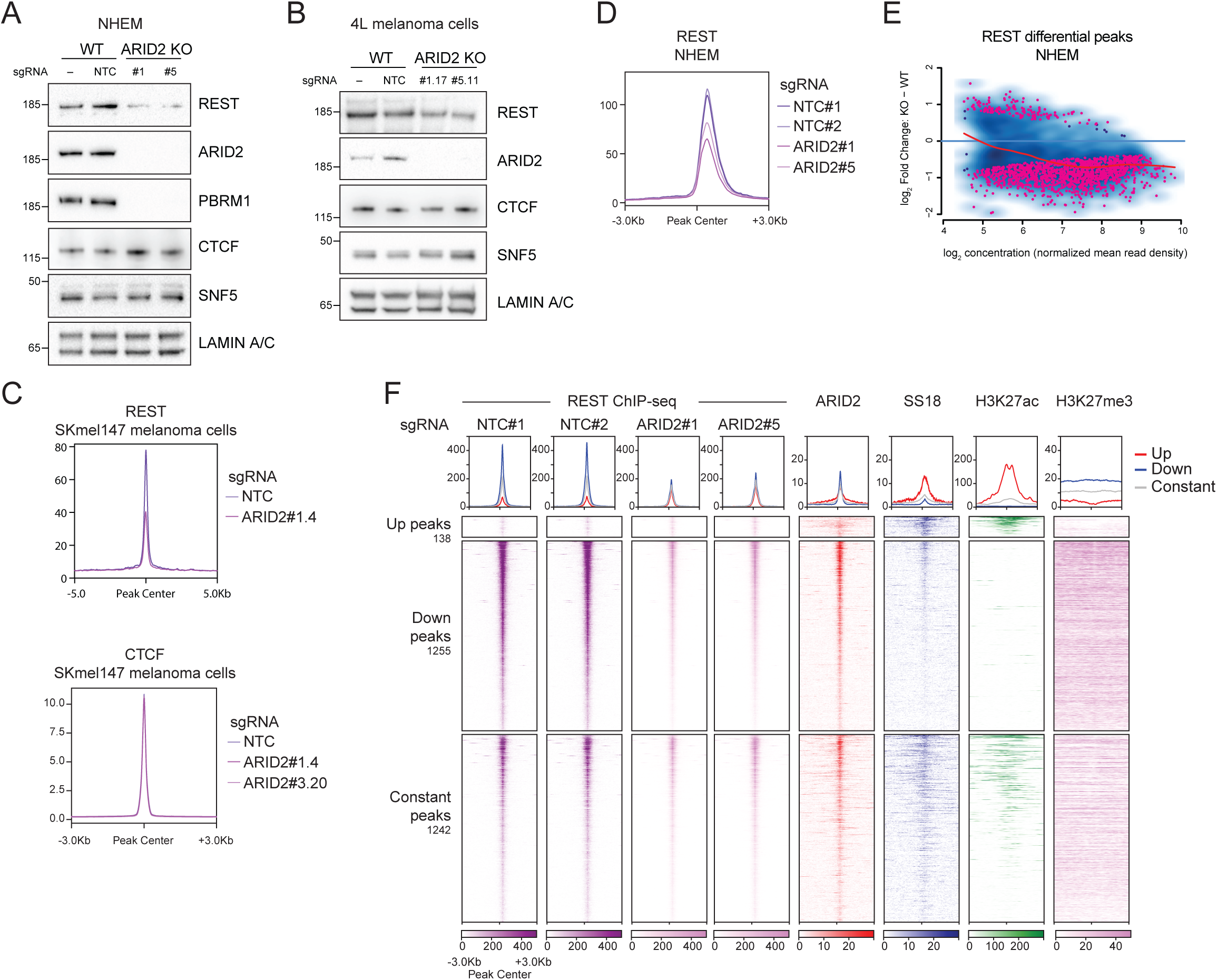
PBAF loss impairs REST binding at repressed chromatin regions. (A-B) Western blot analysis of ARID2 WT and KO chromatin extracts in (A) NHEM and (B) 4L melanoma cells. NTC, non-targeting control sgRNA. LAMIN A/C was used as loading control. (C) Metagene profile of REST and CTCF ChIP-seq in ARID2 WT and KO SKmel147 melanoma cells. (D) Metagene profile of REST ChIP-seq in ARID2 WT and KO NHEMs. (E) MA plot displaying differentially bound REST peaks upon ARID2 KO, as determined by DiffBind analysis. Peaks are plotted according to the extent of the change (fold change KO-WT) and their normalized read density. (F) Heatmap showing REST occupancy in NHEM at differentially bound (Up or Down peaks) or constant REST regions upon ARID2 KO. ChIP-seq signal for ARID2, SS18 and the histone marks H3K27ac and H3K27me3 is also shown at the same regions.

To test this hypothesis, we performed ChIP-seq for these TFs in melanoma cells and melanocytes depleted of ARID2 and observed a widespread decrease in REST occupancy, while CTCF genomic binding remained unaltered (**Fig. 5C, D, S5B**). Statistical analysis revealed that reduction of REST occupancy affects a large fraction of its binding sites (47%; FDR>0.05; |log_2_FC|>0.5) but appears particularly relevant at peaks with high REST binding (**Fig. 5E, F**) as well as peaks localized at repressed regions, which also correspond to the fraction that exhibits the highest ARID2 enrichment (**Fig. 5F**). Constant (|log_2_FC|<0.5) and upregulated regions, the latter of which represent a minor fraction of total sites (0.05%), are generally associated with weaker REST binding and are instead enriched in more active chromatin marked by H3K27ac and SS18. Together, these data demonstrate that REST characterizes a specific set of repressed, ATP-dependent PBAF regions, potentially influencing their sensitivity to PBAF remodeling. In turn, PBAF is required to favor REST access to its target sites, pointing towards a dynamic control mechanism of these repressed regions.

### PBAF loss triggers derepression of REST targets

Despite the unique functional connection between REST and the PBAF complex, this chromatin remodeler is bound to a much larger set of targets than the transcriptional repressor (**Fig. S4A, C**). We therefore sought to investigate whether PBAF-mediated regulation of REST activity impacts gene expression. RNA-seq in WT and ARID2 KO NHEM and melanoma cells revealed different gene expression profiles associated with these distinct cell models (**Fig. 6A, B; Table S4**). While melanoma cells up- and downregulated a similar number of genes upon ARID2 loss (459 and 480, respectively), in NHEM, most of the differentially expressed genes (DEGs) were downregulated (370) and only a handful increased their expression (70). Pathway analysis of these datasets revealed little commonality among decreased genes (**Fig. S6A, B; Table S2**), indicating that some genes controlled by PBAF are context dependent. However, among the upregulated genes, we found that both normal and cancer cells increased the expression of REST targets, as shown by TF enrichment analysis (**Fig. 6C, D; Table S2**). These genes belonged to pathways associated to synapse organization and activity, as well as ion channels, in line with the role of REST as repressor of neuronal genes in non-neuronal cells^41–44^. Importantly, we found these pathways enriched in both NHEM and 4L melanoma cells, and similar terms were found in highly invasive SKmel147 melanoma cells depleted of ARID2 in our previous study^30^.

**Figure 6.**
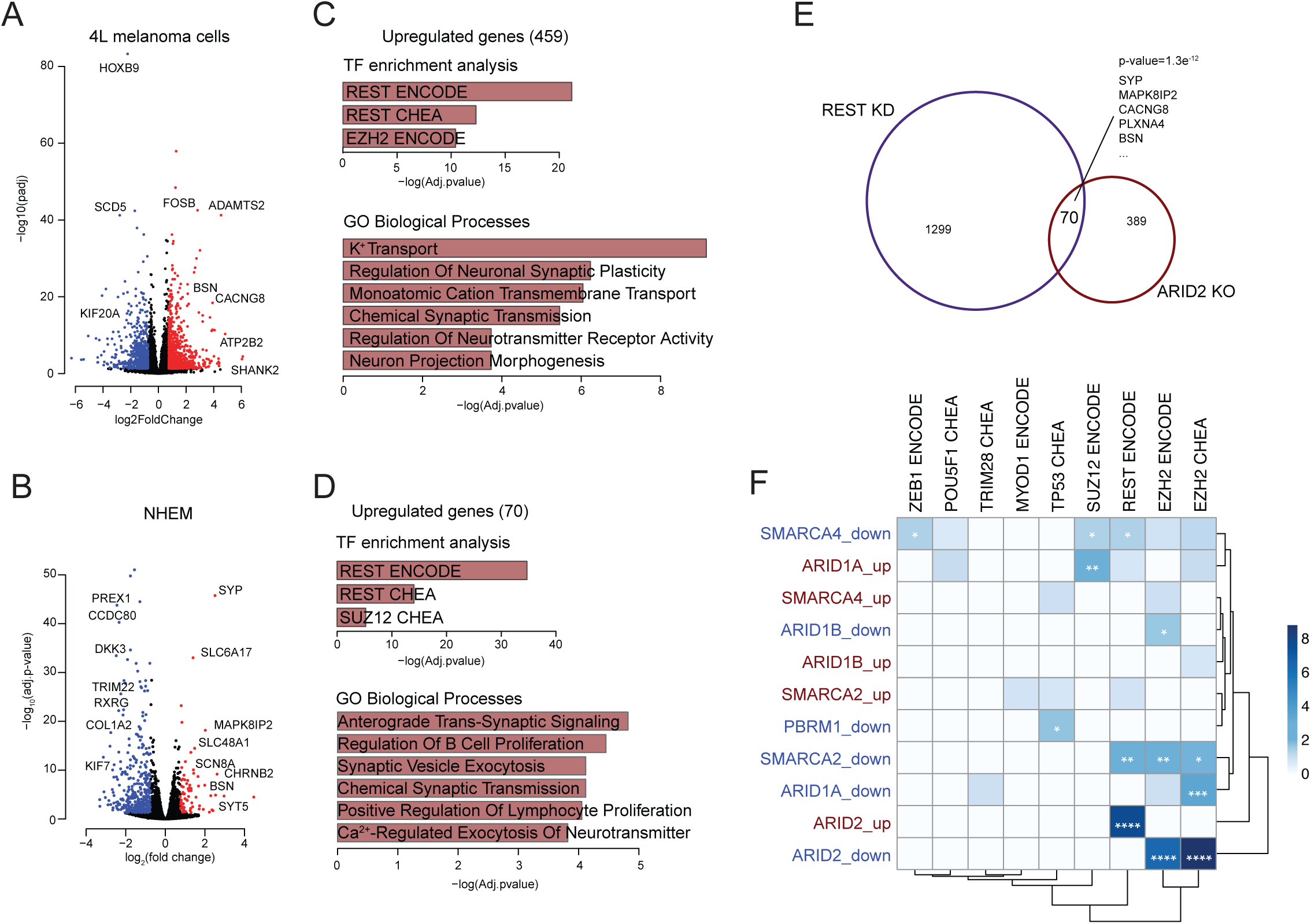
PBAF loss triggers derepression of REST target genes. (A-B) Volcano plots depicting RNA-seq data in ARID2 KO vs. WT (A) 4L melanoma cells and (B) NHEM. (C-D) TF enrichment and Biological Processes GO analyses of genes upregulated (Adj. pvalue <0.05; FC>0.75) upon ARID2 KO in (C) 4L melanoma cells and (D) NHEM. (E) Venn diagram displaying the intersection between upregulated genes in REST KD and ARID2 KO 4L melanoma cells. Overlap significance was calculated by Fisher’s exact test. (F) Heatmap showing TF enrichment GO analysis of genes up or downregulated, as determined by differential expression analysis comparing TCGA SKCM samples mutated for one of the listed SWI/SNF component vs. WT samples. *, pvalue<0.05; **, pvalue<0.01; ***, pvalue<0.001; ****, pvalue< 0.0001.

To further understand the transcriptional program co-regulated by PBAF and REST, we performed RNA-seq in REST knockdown (KD) melanoma cells and compared it with ARID2 KO in the same cell line (**Fig. S6C-D; Table S4**). When we directly compared transcripts upregulated upon ARID2 or REST depletion, we found that the significant overlapping group included neuronal and synaptic genes (**Fig. 6E, S6E**), indicating that PBAF regulation is critical for this specific set of REST targets.

Finally, we sought to extend our analysis to human melanoma patients, given the high rate of ARID2 mutations in this cancer type. We therefore examined public data from the Skin Cutaneous Melanoma (SKCM) TCGA cohort (n=363) and focused on patient samples bearing mutations in SWI/SNF-encoding genes (**Fig. S6F**). We selected the six most highly mutated SWI/SNF components in this cancer type (PBAF-specific ARID2 and PBRM1; BAF-specific ARID1A and ARID1B; core components SMARCA4 and SMARCA2) and compared their gene expression profiles *vs.* their respective WT samples (**Fig. 6F**). Differential expression analysis revealed that downregulated genes in mutant SWI/SNF samples are often enriched in PRC2 targets, suggesting that genetic alterations in SWI/SNF genes lead to PRC2 overactivation, including in ARID2-mutant samples, and in line with the ARID2 KO transcriptome (**Fig. S6A, B**). However, when we focused on upregulated genes, we found that only samples bearing mutations in ARID2 were enriched for REST targets, confirming that even in patients, specific loss of ARID2 function and consequent PBAF impairment reduces the ability of REST to repress its target genes. In conclusion, our data reveals a unique role for PBAF at repressed regions where it facilitates REST binding and silencing activity, a mechanism that is disrupted by ARID2 mutations in melanoma patients.

## Discussion

In recent years, an increasing number of studies have been devoted to identifying and characterizing the biochemical composition, hierarchical assembly, and preference in genomic binding of the SWI/SNF complex subfamilies, yet it remains unclear how the molecular function of these distinct protein assemblies differs. In this study, we elucidated an unappreciated role for the PBAF complex at repressed genomic sites, where its ATPase activity is insufficient to establish active chromatin, but rather is required to facilitate DNA access to the repressive TF REST.

Much of our understanding of the SWI/SNF mechanism of action is based on studies that analyze common and/or catalytic complex subunits, such as the ATPase BRG1. This holds true for the concept that SWI/SNF actively evicts Polycomb complexes^9,15^. To tease apart the specific functions of the PBAF complex, we profiled the epigenomic landscape specifically associated with PBAF regions in melanocytes and melanoma cell models, where this remodeler exerts an important role^30^. Notably, we found that a subset of PBAF sites was enriched for PRC2 and the H3K27me3 mark that it deposits, challenging the notion that all SWI/SNF subcomplexes are capable of opposing PRC1/2 and restoring active chromatin with the same efficiency. Interestingly, melanocytes demonstrated a higher enrichment for these heterochromatic marks than melanoma cells, possibly reflecting the more condensed chromatin structure characteristic of differentiated cells compared to plastic cancer cells^34,45^. Consistent with our findings, it was recently reported that cBAF, but not PBAF, could remove Polycomb-associated marks when recruited to the bivalent *Nkx2.9* locus in mESCs^46^, a finding consistent with a lack of PBAF-PRC2 opposition. Moreover, it suggests that our findings are not restricted to the melanocytic lineage.

PBAF-specific regions displayed lower chromatin accessibility compared to cBAF/ncBAF-specific or shared sites, suggesting that PBAF remodeling activity is insufficient to properly open its target regions in the absence of other SWI/SNF complexes. Using BRM014 to block SWI/SNF ATPase activity, we found that only a small fraction of PBAF sites responded with reduction in accessibility; conversely, cBAF/ncBAF regions were largely sensitive to BRM014, quickly losing accessible chromatin. Therefore, PBAF displays distinct remodeling dynamics compared to other SWI/SNF subcomplexes. One possibility for this difference in complex behavior is that PBAF’s intrinsic structural features limit its ability to efficiently open chromatin in absence of other remodelers *in vivo*. Recent cryoEM studies that solved BAF and PBAF structures have revealed that these two complexes adopt different conformations to engage the nucleosome^3,5,47^; yet further studies are needed to understand how these distinct characteristics affect remodeling *in vivo*. Another possibility is that the chromatin features of PBAF-bound sites dictate their poor susceptibility to ATPase-mediated remodeling, as suggested to explain different sensitivities to SWI/SNF chemical inhibition^13,14^.

We speculate that the presence of complex-specific TFs could also contribute to influence remodeling sensitivity at their target regions. In fact, recent evidence indicates that the SWI/SNF complex probes the genome, transiently colocalizing with both active and repressive chromatin marks, and it productively engages and mobilizes nucleosomes only in association with specific TFs^8,9^. Therefore, we inquired whether different TFs were linked to distinct responses to SWI/SNF inhibition. We found that BAF- and PBAF-unique regions are enriched in different sets of TFs. While BAF sites are enriched in neural crest-specific activating TFs found at promoters and enhancers, such as MITF and SOX10^48,49^, PBAF regions present unique binding for the insulator protein CTCF and the repressive TF REST. Interestingly, both TFs recognize a large binding motifs of >17 base pairs with high specificity and mediate DNA demethylation^10,50^; yet they respond differently to remodeling activity. CTCF and REST have been shown to characterize distinctive clusters of genomic regions that respectively increase or slowly decrease chromatin accessibility levels in response to SWI/SNF inhibitors in mESCs^12^. We now demonstrate that both CTCF- and REST-enriched clusters are exclusively associated with PBAF regions with unique responses to SWI/SNF inhibition. Importantly, our data suggest caution when using these TFs to generally inform on SWI/SNF function, as they prevalently associate with PBAF, at least in the melanocytic lineage. Moreover, it has been shown that the Imitation Switch (ISWI) family of remodelers is primarily responsible for CTCF binding and establishment of local accessibility^10,51^. Further studies are therefore needed to clarify whether multiple chromatin remodelers work together at CTCF sites. In addition, we expect different TF-remodeler associations to be found in distinct tissues and/or organisms. By analyzing ChIP-seq data profiling SWI/SNF complexes in mouse mammary epithelial cells^13^, we found the PBAF complex associated with the TF Ronin/GFY, which is also characterized by a large binding motif, suggesting that a unique binding ‘grammar’ might be linked to PBAF occupancy.

REST is a silencing TF that we previously linked to PBAF-enriched regions in melanoma cells^30^. Here we found that this transcriptional repressor characterizes ATPase-sensitive PBAF-specific sites that are enriched in repressive marks. Notably, while CTCF chromatin binding is not affected by ARID2 depletion and consequent PBAF disassembly, REST occupancy is reduced in absence of PBAF, in particular at repressed regions where it colocalizes with this remodeler. REST is a lineage-specific TF that recruits histone deacetylases and methyltransferases to repress neuron-specific genes involved in synaptogenesis, axon guidance and synaptic structure in all non-neuronal cells^42,52^, but its function is particularly relevant in neural crest cells (NCCs), as they differentiate into many cell types, including neurons and melanocytes^53,54^. We propose that PBAF’s ability to establish limited levels of accessibility at relevant developmental genes is required to enable the binding of silencing factors, such as REST, that in turn ensure a repressive state by recruiting their associated corepressors. In fact, RNA-seq of ARID2 KO melanoma cells and melanocytes revealed the upregulation of a common REST gene expression signature. This demonstrates that PBAF deficiency prevents REST function at its target genes, which in turn results in derepression of neuronal genes. Given the importance of these genes in defining neuronal functions and the observed pigmentation defects in NCC-specific *Rest* conditional KO (cKO) models^54^, we speculate that impairment of PBAF-mediated regulation of REST targets might interfere with proper NCC development and be linked to neurocristopathies. Consistently, PBAF is known to control NCC formation along with CDH7, whose dysfunction causes CHARGE syndrome^55,56^. However, PBAF function could also be relevant in cancer, as it has been shown that melanoma tumors can form synaptic-like connections with surrounding cells, such as keratinocytes and nerves, to promote tumor growth, angiogenesis and dissemination^57,58^. In keeping with this idea, melanoma patient samples bearing ARID2 mutations upregulate REST target genes, indicating that ARID2 alterations may predict for this synaptic signature. Future studies will be aimed at determining the functional consequences of this conserved signature and whether it plays a role in promoting melanoma progression and/or allowing adaptation to certain host microenvironments.

In conclusion, our study provides insight into the relationship between distinct SWI/SNF remodelers and TFs, whose tissue-specific function might help to explain why genetic alterations in individual SWI/SNF components are enriched in particular tumor types.

## Acknowledgments

The authors thank the laboratory of Roland Friedel for expertise and advice; Ana Hahn and Kevin Mohammed for NGS support; Austin Meadows and Nivetha Aravind for technical support; Ivan Raimondi for advice; the Bernstein lab for discussions and feedback; the Bioinformatics for Next Generation Sequencing (BiNGS) core at the Icahn School of Medicine at Mount Sinai (ISMMS) for bioinformatic support, and the Flow Cytometry core at ISMMS for support with cell sorting. This study was supported by American-Italian Cancer Foundation and National Cancer Center and SBDRC Transition to Independence Minigrant (funded through NIAMS/NIH SBDRC P30 AR079200-02) to E.G., NIH/NCI F30 CA253988-02 to C.B.N.; American Skin Association to D.F.; and NIH/NCI R01CA154683, MRA Pilot Award for Women in Melanoma Research, and MRF Established Investigator Award to E.B. This work was supported in part by Tisch Cancer Institute of the ISMMS Cancer Center, support grant P30CA196521; Scientific Computing supported by the Office of Research Infrastructure of the NIH under award no. S10OD026880 to ISMMS; and the ISMMS Genomics Technology Facility.

## Author contributions

Conceptualization, E.G. and E.B.; investigation, E.G., C.B.N., S.M., V.K.C.; formal analysis, E.G., C.B.N., S.C., D.H.; writing – original draft, E.G. and E.B; writing – review & editing, E.G., C.B.N., S.C., S.M., V.K.C., D.F., D.H. and E.B.; visualization, E.G.; expertise and methods, D.F.; data curation, E.G.; supervision, D.H. and E.B.; funding acquisition, E.G., C.B.N., and E.B.

## Supplemental information

Document S1. Figures S1–S6

Table S1. PBAF-, shared and BAF-enriched regions, related to Fig. 1

Table S2. Complete GO analysis, related to Fig. 2 (tab 1), 6 (tab 2, 3) and S6 (tab 4)

Table S3. Motif analysis, related to Fig. 2 (tab 1) and S3 (tab 2)

Table S4. DEGs in RNA-seq analysis, related to Fig. 6 (tab 1, 2, 4) and S6 (tab 3)

## STAR Methods

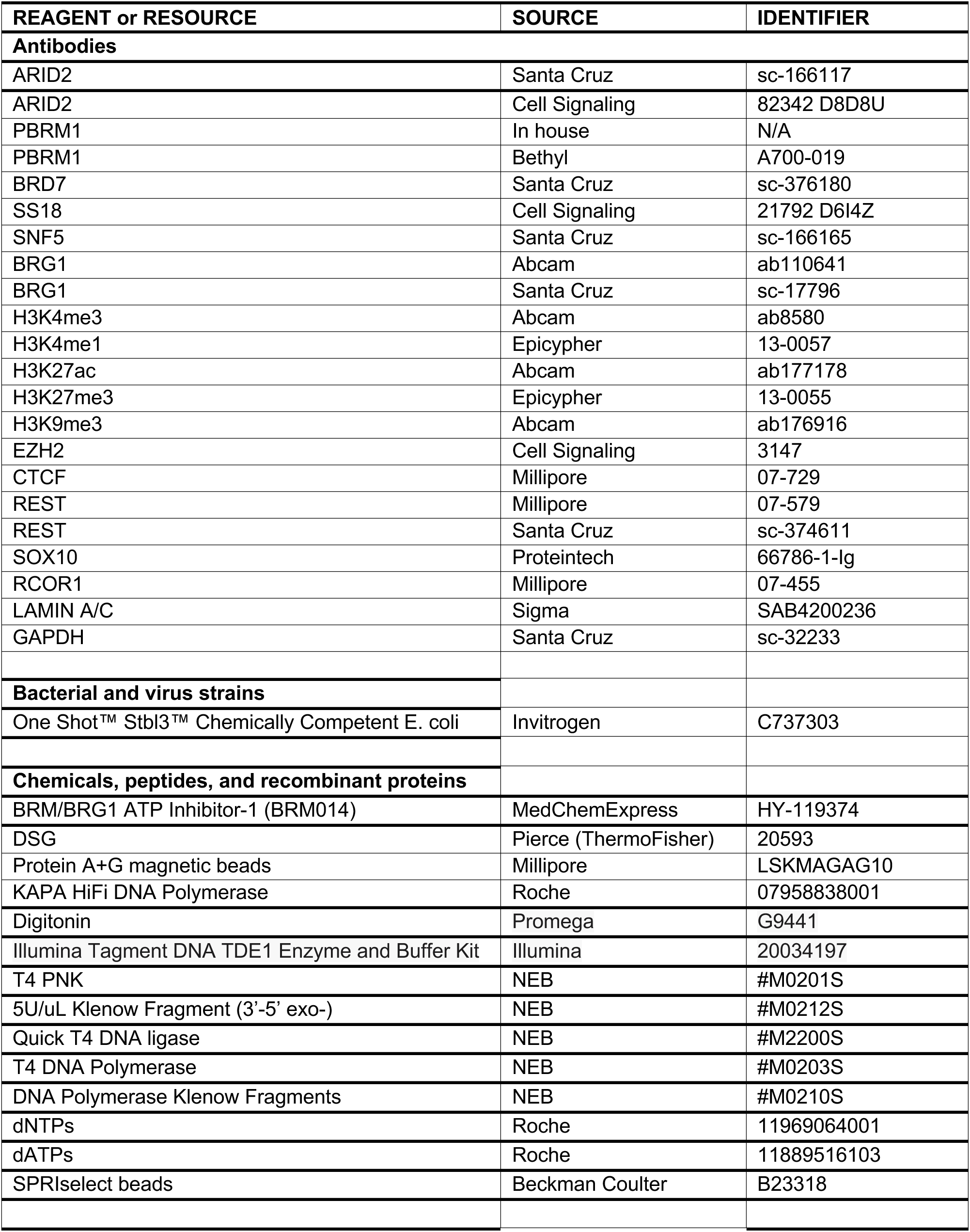
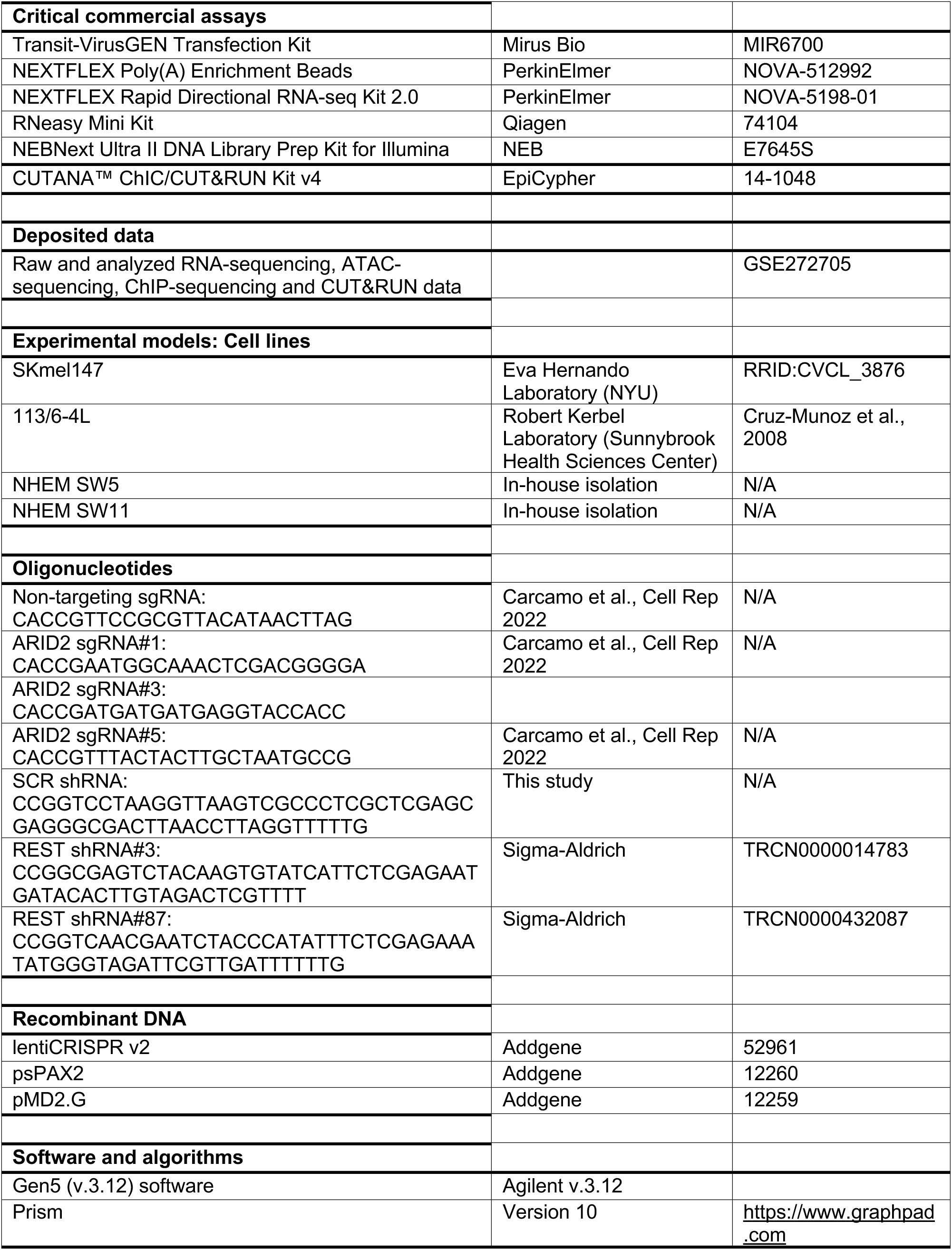
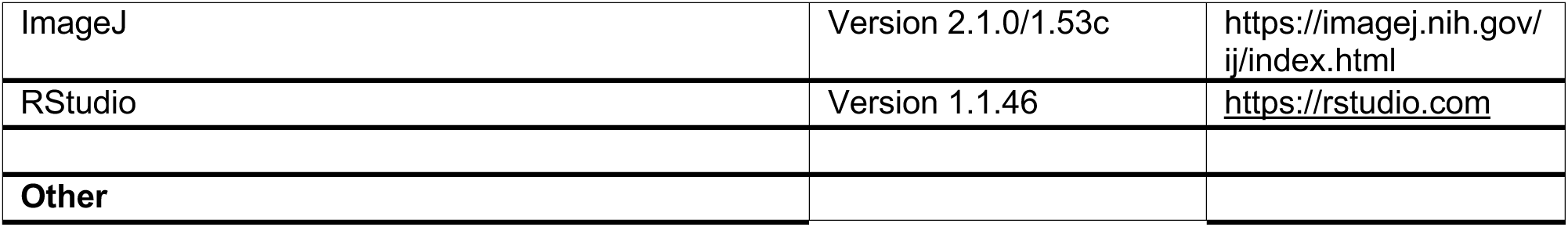

### RESOURCE AVAILABILITY

#### Lead contact

Further information and requests for resources and reagents should be directed to and will be fulfilled by the lead contact, Emily Bernstein.

#### Materials availability

Reagents used in this study are publicly available or available from the lead contact upon request.

#### Data and code availability

Raw and processed data from RNA-seq, ATAC-seq, and ChIP-seq experiments generated in the current study have been deposited in the Gene Expression Omnibus (GEO) repository under the accession number GSE272705. Published ChIP-seq datasets generated in SKmel147 melanoma cell lines and in non-tumoral murine models are available through the GEO repository (accession numbers GSE172383 and GSE226953, respectively). TCGA melanoma data are publicly available in the NCI Genomic Data Commons (GDC) data portal under project ID TCGA SKCM (https://portal.gdc.cancer.gov/projects/TCGA-SKCM). This paper does not report original code.

### EXPERIMENTAL MODELS

#### Cell culture

Melanoma cell lines SKmel147 and 113/6-4L were cultured in DMEM supplemented with 10% FBS (R&D System) and 1X penicillin-streptomycin (Gibco). HEK293T cells were used for virus production and were maintained in DMEM supplemented with 10% FBS and 1X penicillin-streptomycin. Primary normal human epithelial melanocytes or NHEM (batches SW5-SW11) were extracted and cultured as described in^59^. Briefly, neonatal foreskins were collected in PBS supplemented with 1X Antibiotic-Antimycotic (Gibco #15240-062), and stored at 4C for no longer than 48 hr. Skins were then washed once in 70% EtOH and 3x in PBS supplemented with 1X Antibiotic-Antimycotic, cut into 3-4 pieces in HBSS (Gibco #24020-117) and placed in Dispase II solution (Roche #04942078001) at 4C O/N. Epidermis was then separated from the dermis in HBSS, floated in 1 ml of 0.25% trypsin (Gibco #15400-054), and incubated in 37C for 3 min. Detached cells were then collected in 4 mL of DMEM with 10% FBS and spun at 1200 rpm for 5 min. The last step was repeated a second time and the two cell pellets were consolidated and resuspended in Melanocyte Growth Media 254 supplemented with Human Melanocyte Growth Supplement (HMGS, Life Technologies #S-016-5) and 0.2 mM CaCl2, for 3-5 days. Cells were then rinsed in PBS, 1:100 Anti-Anti, and trypsinized in 1 ml 0.05% Trypsin for exactly 3 min to separate melanocytes from fibroblasts and keratinocytes. Cells were then collected into 4 mL of DMEM with 10% FBS, spun at 1200 rpm for 5 min and resuspended and cultured in Melanocyte Growth Media 254 supplemented with HMGS, 0.2 mM CaCl2, and 10 ng/ml PMA (Sigma #P8139).

## METHOD DETAILS

### Plasmid and infections

For CRISPR-mediated knockout in melanoma cells and primary melanocytes, gRNAs targeting *ARID2* and well as non-targeting control (NTC) gRNAs were cloned into the lentiCRISPRv2 (Addgene #52961). Fore REST knockdown experiments, shRNAs constructs provided in a pLKO.1 lentiviral backbone were obtained from the RNAi Consortium (TRC, Open Biosystems). Scrambled (SCR) shRNA was used as control. For gRNA and shRNA sequences, see Table S4. To produce lentiviral particles, HEK293T cells were seeded in 10 cm tissue culture dishes and co-transfected with 5 μg of lentiviral expression constructs, 3.75 μg of psPAX2 and 1.25 μg pMD2.G vectors using the TransIT-VirusGen (Mirus Bio) transfection reagent diluted in Opti-MEM medium (Gibco), following manufacturer’s recommendations. 48 hours post-transfection, supernatants were collected and passed through a 0.45 μm PVDF filter to remove cells and debris. Upon infection, cells were selected with puromycin (1 μg/ml). For CRISPR-mediated knockout in melanoma cells, limiting dilutions were used to generate colonies from single cells. Individual colonies were isolated and further expanded prior to screening by immunoblot. For primary melanocytes, infected cells were kept as pool, without isolating single clones. KO cell lines are identified by the sgRNA used followed by the clone number, when single clones were isolated (e.g., KO#1.17).

### Cell growth assay

To test cell growth upon BRM014 treatment, 2×10^3^ 4L melanoma cells were plated into black polystyrene 96-well plates with clear flat bottom (Agilent) in triplicate. BRM014 inhibitor was added 24h post plating. Plates were imaged on brightfield every 6h in a 48h time course with a 4X PL FL magnification in a Cytation 7 automated microscope (Agilent). Cell count was performed using the Gen5 (v.3.12) software (Agilent), normalizing each measurement to time 0. Growth curves were plotted on GraphPad Prism version 10 (GraphPad Software) using normalized cell counts.

### Whole-cell protein extraction and chromatin fractionation

Chromatin fractionation was performed as described^60^. Briefly, cell pellets were lysed on ice for 8 min in Buffer A (10 mM HEPES pH 7.9, 10 mM KCl, 1.5 mM MgCl2, 0.34 M Sucrose, 10% glycerol supplemented with 1:200 protease inhibitors cocktail and 1 mM DTT) + 0.1% Triton X100. Cell extracts were centrifuged for 5 min at 1850 g and the pellets were washed with 1 ml of Buffer A (supplemented with 1:200 protease inhibitors cocktail and 1 mM DTT). Samples were spun down for 5 min at 1850 g and the pellets were resuspended in No Salt Buffer (3 mM EDTA, 0.2 mM EGTA supplemented with 1:200 protease inhibitors cocktail and 1 mM DTT) and incubated on ice for 30 min with occasional mixing. Nuclear extracts were centrifuged for 5 min at 1850 g and chromatin pellets were resuspended in 200 μl Buffer A (supplemented with 1:200 protease inhibitors cocktail and 1:200 Benzonase). Pellets were solubilized for 15 min at 37C shaking and subsequently used for Immunoblot. For whole-cell protein extraction, cells were lysed in RIPA lysis buffer (50mM Tris-HCl pH8, 150mM NaCl, 1% NP-40, 0.5% Sodium deoxycholate, 0.1% SDS supplemented with protease inhibitors cocktail and 1:500 Benzonase) and incubated on ice for 15 minutes with occasional mixing. Lysates were sonicated on high level, 5 cycles 30s ON, 30s OFF and centrifuged at max speed for 10 minutes. Protein concentrations were quantified using Bicinchoninic Acid (BCA) assay (Pierce), according to manufacturer’s instructions.

### Immunoprecipitation (IP) with chromatin extracts

Chromatin extracts were prepared as mentioned above using 10-20×10^6^ cells per IP. After BCA quantification, chromatin extracts were diluted in Buffer C (50 mM Tris HCl pH 7.5, 150 mM NaCl, 2mM EDTA, 0.05% NP-40) to equalize protein amounts (*e.g.* 500 μg per condition). Diluted extracts were pre-cleared with 20 μl of Magna ChIP Protein A+G magnetic beads (Millipore) for 1 hours in rotation at 4C and 5% of extract was saved as input. Precleared extracts were then incubated overnight with antibodies (see Table S4), followed by 2 h incubation with 25 μl of magnetic beads. Beads were subsequently washed once with Buffer G150 (50 mM Tris HCl pH 7.5, 150 mM NaCl, 0.5% NP-40), twice with Buffer G250 (50 mM Tris HCl pH 7.5, 250 mM NaCl, 0.5% NP-40) and once with TE buffer and proteins were eluted in 50 μl of 4X Laemmli loading buffer (Bio-Rad) and boiled 10 minutes at 95C.

### Glycerol gradient sedimentation

Glycerol gradient sedimentation was performed according to^61^. 40-50 million melanoma SKmel147 cells were collected, lysed in Nuclear isolation buffer (20 mM HEPES pH 7.9, 25 mM KCl, 10% glycerol, 5 mM MgCl2, 0.05 mM EDTA, 0.1% NP-40, 100 nM PMSF, supplemented with protease inhibitors) and washed once with the same buffer without NP-40. The nuclear extracts were then resuspended in Nuclear lysis buffer (10 mM HEPES pH 7.6, 3 mM MgCl2, 100 mM KCl, 0.1 mM EDTA, 10% glycerol). 300 mg/mL of ammonium sulfate powder was added to the mixture and incubated on ice for 20 min. Proteins were pelleted by ultracentrifugation at 150,000 g for 30 minutes and resuspended in 100 mL of HEG1000 buffer (25 mM HEPES pH 7.6, 0.1 mM EDTA, 12.5 mM MgCl2, 100 mM KCl, supplemented with protease inhibitors). A glycerol gradient (10-30%) was prepared using HEG1000 buffer without glycerol and with 30% glycerol. Next, the protein lysates were layered on top of the 10-30% glycerol gradients (10 mL) and fractionated by centrifugation at 40,000 rpm for 16 h using a SW32Ti rotor (Beckman Coulter). Fractions (20x) of 500 mL were collected sequentially from the top of the gradient, diluted with Laemmli loading buffer and boiled for 10 minutes before being used for immunoblotting.

### Immunoblotting

Protein lysates were mixed with 2X or 4X Laemmli loading buffer supplemented with 2-mercaptoethanol and boiled at 95C for 5 min prior to immunoblotting. Samples were run on a NuPAGE Bis-Tris 4-12% precast gels (Invitrogen) and wet transferred onto a PVDF Immobilon transfer membrane (Millipore) for 1 hour at 30V using Bis-Tris transfer buffer with 20% methanol for high molecular weight targets. Membranes were first incubated with Amido Black to verify uniform loading and incubated with blocking buffer (5% Blotting-grade Blocker (Bio-Rad) in Tris-Buffered Saline, 0.1% Tween (TBS-T)) for 1 hour, followed by overnight incubation with the respective primary antibodies (listed in Table S4). The next day membranes were incubated with the respective secondary antibodies, rinsed with TBS-T, and developed using Clarity Western ECL Substrate (Bio-Rad).

### Chromatin Immunoprecipitation (ChIP-seq)

ChIP samples were processed as previously described^30,59^ with modifications. To detect histone modifications, 10-15×10^6^ cells per condition were crosslinked directly on plate with 1% Formaldehyde for 10 minutes at room temperature. For chromatin remodeler subunits and transcription factors, 10-40×10^6^ cells were double cross-linked with 0.25 M disuccinimidyl glutarate (DSG), followed by 1% formaldehyde. Single and double crosslink reactions were quenched with 0.125 M glycine for 5 minutes at room temperature, cells were thoroughly washed with PBS and scraped off the plates in 1 ml ice-cold PBS. Chromatin was then aliquoted and pelleted at up to 1500 g for double crosslinked cells at 4C and stored at −80C or used directly for ChIP. Nuclei were isolated by resuspending cell pellets in 10ml of Cell Lysis Buffer (10mM Tris pH 8.0, 10mM NaCl, 0.2% NP-40 supplemented with 1:200 protease inhibitors cocktail and 1:2000 DTT) rotating for 10 min at 4C and spun down at 2000g for 10 min at 4C. 15. Nuclear pellets were then resuspended (400μL/10M cells) in Nuclear Lysis Buffer (50mM Tris pH 8.0, 10mM EDTA pH 8.0, 1% SDS supplemented with 1:200 protease inhibitors cocktail and 1:2000 DTT) and incubated on ice with frequent mixing. Cells were sonicated for 20-30 cycles, 30 secs on, 30 secs off, at low (single crosslink) or high (double crosslink) intensity in a Bioruptor sonicator (Diagenode). After sonication, samples were centrifuged at max speed for 10 minutes at 4C, and the supernatant-containing chromatin was diluted 1:4 with IP Dilution buffer (20 mM Tris pH 8, 2 mM EDTA, 150 mM NaCl, 1% Triton-X, 0.01% SDS, 100 nM PMSF, supplemented with protease inhibitors). Where indicated, Drosophila spike-in chromatin was added to the diluted chromatin, followed by pre-clearing with Magna ChIP Protein A+G magnetic beads (Millipore) for 2 hours rotating at 4C. 50 μl of chromatin was saved for input control and chromatin extracts were incubated with the designated antibody O/N in rotation at 4C (ChIP-seq antibodies are listed in Table S4). The following day, 50 μl of magnetic beads per condition were washed and incubated with the antibody-containing extract for 2 hours rotating at 4C. Beads were then washed once with cold IP Wash I buffer (20 mM Tris pH 8, 2 mM EDTA, 50 mM NaCl, 1% Triton-X, 0.1% SDS, supplemented with protease inhibitors cocktail), twice with cold High Salt buffer (20 mM Tris pH 8, 2 mM EDTA, 500 mM NaCl, 1% Triton-X, 0.01% SDS, supplemented with protease inhibitors cocktail), once with cold IP Wash II buffer (10 mM Tris pH 8, 1 mM EDTA, 0.25 LiCl, 1% NP-40, 1% deoxycholic acid, supplemented with protease inhibitors cocktail) and twice with cold TE buffer (5 mM Tris pH 7.4, 1 mM EDTA). DNA was eluted and reverse crosslinked at 65C with mixing in 50 μl of Elution Buffer (300mM NaCl, 1% SDS, 100mM NaHCO3) + 2 μl RNAse A for 2 hours, followed by overnight incubation at 65C with 2.5 μl of Proteinase K. DNA was isolated by 2.5X of SPRI beads (Beckman Coulter), following manufacturer’s instructions. Library preparation for multiplexes sequencing was carried out according to Illumina’s recommendations. 1-8ng of DNA input or ChIP DNA was end-repaired with T4 DNA polymerase (NEB), DNA polymerase I, large (Klenow) fragment (NEB) and T4 polynucleotide kinase (NEB) in the presence of deoxyribonucleotide triphosphates (dNTPs) (Roche). Next, dA-tailing was done using Klenow fragment (3′→5′ exo-, NEB) with deoxyadenosine triphosphate (dATP) (Roche) in NEB Buffer 2. Barcoded Illumina Truseq adaptors were ligated using Quick Ligase (NEB). Libraries were amplified using KAPA HiFi DNA Polymerase (KAPA Biosystems), with optimal number of cycles determined by qPCR, and size selected using SPRI beads. Resulting libraries were quantified by Qubit (Thermo Fisher) and average DNA fragment size was calculated using an Agilent bioanalyzer or Tape Station system. Libraries were then multiplexed and sequenced on a NextSeq500/550 or NExtSeq2000 instrument (Illumina) at 75bp single-end or paired-end reads.

### Cleavage Under Targets & Release Using Nuclease (CUT&RUN)

CUT&RUN was performed using CUTANA ChIC/CUT&RUN kit (Epicypher #14-1048) following manufacturer’s protocol. Briefly, 5 × 10^5^ cells were coupled with Concanavalin A beads, permeabilized with 0.01% Digitonin and incubated overnight at 4C with 0.5 μg target antibody in antibody buffer (antibodies listed in Table S4). The following day, cells were first incubated for 10 minutes with micrococcal nuclease fused to proteins A and G (pAG-MNase), which was then activated by CaCl2 addition. After 2h incubation at 4C, the reaction was stopped with stop buffer and E. coli DNA was added as spike-in DNA. The DNA was then isolated using SPRI beads and quantified with Qubit. Libraries were prepared using NEBNext Ultra II DNA library kit, following manufacturer’s recommendations. Fragment size was detected using a Tape Station system and multiplexed libraries were sequenced on a NExtSeq2000 instrument (Illumina) at 50bp paired end reads.

### Assay for Transposase-Accessible Chromatin (ATAC-seq)

The Omni-ATAC protocol was used to profile chromatin accessibility^62^ with minor modifications. 5×10^4^ cells were harvested, treated with Digitonin (Promega) and tagmented with 2.5 μl of TDE1 Enzyme (Illumina). The correct number of amplification cycles was determined by qPCR and resulting libraries were isolated and size selected using SPRI beads. Libraries were quantified by Qubit and average DNA fragment size was determined using an Agilent bioanalyzer or Tape Station system. Multiplexed libraries were then sequenced on Illumina NextSeq2000 at 50bp paired-end reads. For ATAC-seq upon ATPase inhibitor treatment, samples were treated with 1 μM of BRM014 compound (MedChemExpress) starting the treatment at different time points, such that all the samples could be collected at the same time.

### RNA-seq

Total RNA was extracted from about 2-4×10^6^ cells using the RNeasy Mini Kit (Qiagen) according to manufacturer’s protocol, including DNase digestion. 1-2 μg of total RNA was used to isolate poly-A RNA with NEXTFLEX Poly(A) beads 2.0 (Perkin Elmer), while libraries were prepared using the NEXTFLEX Rapid Directional RNA-seq Kit 2.0 (PerkinElmer). Samples were quantified by Qubit and quality was assessed by Agilent bioanalyzer or Tape Station system. Multiplexed libraries were sequenced on a NextSeq500/550 in a 75bp single-end format.

### ChIP-seq and CUT&RUN analysis

Adapters were trimmed using Trimmomatic (v.0.36)^63^ or NGmerge (v.0.3)^64^ for single- or paired-end reads, respectively. Reads were aligned to the human reference genome (GRCh38/hg38) using Bowtie2 (v.2.4.1)^65^ with standard parameters, and reads quality was assessed using fastQC. samtools (v.1.9)^66^ was used to generate bam files and filter out low-quality (Phred quality score <30), mitochondrial and duplicated reads. Significant binding peaks were determined using macs2 v2.1.0^67^ for narrow peaks, and SICER 2.0^68^ for broad peaks, with matching input files as control. Cut-off values (q-values) were determined post-hoc, testing several q-values based on signal-to-background ratio and using macs2 --cutoff-analysis parameter for additional guidance. Peaks located in ENCODE blacklisted regions were excluded. Coverage tracks (bigwig files) were generated from filtered bam files using deepTools (v. 3.5.1)^69^ bamCoverage with parameters –normalizeUsingRPGC –binsize 10. Metagene plots and heatmaps showing signal enrichment over genomic regions were generated by first calculating intensity scores for each genomic site using deepTools computeMatrix, and then plotting heatmaps and/or metagene plots with plotHeatmap, selecting the center of the peaks as reference point. To remove non-specific signal from accessible chromatin in EZH2 CUT&RUN, ATAC signal was subtracted using deeptools bigwigCompare with the following parameters: --operation ratio --scaleFactors 1:0.01. To precisely quantify H3K27me3 read intensity at SWI/SNF and EZH2 regions, the matrix containing scores per genomic regions calculated with computeMatrix was exported and plotted in R studio. Significance was calculated by Mann-Whitney U test using the wilcox.test function in R.

To identify PBAF-enriched, shared and BAF-enriched regions, ARID2 and SS18 peaks in 4L and SKmel147 melanoma cells and SW11 melanocytes were called using macs2 and compared using bedtools (v.2.29.2)^70^ intersect. The regions are listed in Table S1. The CHIPSeeker 1.34.1 package^71^ was used to annotate peaks and to determine their feature distribution. Promoters were defined as ±1Kb from TSS using the annotation package TxDb.Hsapiens.UCSC.hg38.knownGene. Chromatin state modeling was performed using the chromatin Hidden Markov Model (chromHMM) software as previously described^33,72^. The overlap between each state and SWI/SNF peaks was calculated using bedtools intersect.

For differential binding analysis, bam files generated in ARID2 WT and KO samples were analyzed along with matching input files using DiffBind (v. 3.4.11)^73^, following the package documentation (http://bioconductor.org/packages/release/bioc/vignettes/DiffBind/inst/doc/DiffBind.pdf). Briefly, a binding matrix was generated with dba.counts (summits=250). Drosophila spike-in data were used for normalization when available, otherwise count normalization was performed using the DBA_NORM_NATIVE parameter with background=T. DESeq2 was used to perform differential peak analysis (FDR<0.05 & |log_2_FC|>0.5) and generate MA plots. For visualization purposes, the bam files of WT and KO samples were merged into a master file using samtools merge. Significant peaks were called on this merged file using macs2 with matching input as control and individual coverage tracks were normalized by using DESeq2 scaling factors. Scaled tracks were then plotted with deeptools plotHeatmap to show change in binding at merged peaks.

### ATAC-seq analysis

Adapter trimming, alignment, read filtering and genome coverage calculation were performed as in ChIP-seq and CUT&RUN. ATAC-seq bam files were merged, and peaks were identified as in^12^, by first converting the paired-end bam files to single-end bed format and then running macs2 callpeak function with --shift −100 --extsize 200 parameters. Peaks located in ENCODE blacklisted regions were excluded. For analysis of differentially accessible regions upon BRM014 treatment, two biological replicates for each time point were performed. DiffBind was employed to compare treated samples to the DMSO control sample, as for ChIP-seq data but using the summit=100 parameter to generate the count matrix. ATAC-seq peaks were considered gained or reduced if their accessibility levels were affected at any time point, as calculated by DESeq2 (FDR<0.05 & |log_2_FC|>1). Constant and changing ATAC-peaks were associated to SWI/SNF regions (Table S1) using bedtools window (-w 500). Metagene plots showing change in chromatin accessibility at different SWI/SNF regions were generated by using deepStats (v. 0.3.1 - https://doi.org/10.5281/zenodo.3336593) dsCompareCurves and bigwig files normalized according to DESeq2 scaling factors. Heatmaps depicting changes in ATAC-seq peaks over time were generated with ggplot2. Regions were grouped by k-means clustering in R (v. 4.2.2) by the k-means function with parameter nstart = 10 and using the silhouette function to determine the optimal number of clusters. For motif enrichment analysis of ATAC-seq clusters, de novo motifs were identified using Homer findMotifsGenome.pl (-size 200) with background peaks consisting in the totality of ATAC-seq peaks retrieved (Table S3)

To quantify ATAC-seq read intensity at SWI/SNF regions in parental melanocytes and melanoma cells, the matrix containing scores per genomic regions calculated with computeMatrix was exported and plotted in R studio. Significance was calculated by Mann-Whitney U test using the wilcox.test function in R. For ATAC-SWI/SNF correlation analysis, PBAF and BAF peaks were selected to include only regions shared by fully formed complexes (i.e. ARID2, PBRM1 and BRG1 for PBAF; SS18 and BRG1 for BAF). Deeptools multiBigwigSummary was employed to generate an enrichment matrix over the filtered PBAF or BAF regions and plotCorrelation function was used to visualize plots and compute Spearman correlation.

### RNA-seq analysis

Single-end 75-bp reads were aligned to the human reference genome (hg38/GRCh38) with STAR (v. 2.7.5b)^74^ using the parameters --runMode alignReads -- sjdbOverhang 100 --outFil-terMultimapNmax 10 --outFilterMismatchNmax 10 -- outFilterType BySJout --outFilterIntronMotifs Remove-NoncanonicalUnannotated. Next, the featureCounts function of the Rsubread R package (v. 3.19)^75^ was used to assign reads to genes. Assigned reads were then normalized and differential expression analysis was performed using the R package DESeq2 (v. 1.38.3)^76^. Genes were considered expressed if the sum of raw counts across all samples was >10 for any given gene. Differentially expressed genes were called using adjusted P-value of ≤0.05 and log_2_FC of ≥0.75 or ≤−0.75. Volcano plots were generated with the ggplot2 R package to depict DEseq2 results. Gene Ontology (GO) enrichment analysis was performed on significantly up or downregulated genes using the Enrichr web server^77^ and selecting all expressed genes as background gene set. Transcription factor enrichment analysis and GO Biological Processes categories were selected and adjusted p-values for the top terms were shown as bar plots. The complete list of enriched terms is reported in Table S2.

### TCGA analysis

Processed data from The Cancer Genome Atlas (TCGA - v. 28) were obtained from the National Cancer Institute’s Genomic Data Commons (GDC) Data Portal. The skin cutaneous melanoma (SKCM) raw RNA-seq reads and mutational calls were downloaded using the TCGAbiolinks (v.2.25.3) package^78^. Sample normalization of raw counts was carried out using the median-ratios normalization method from DESeq2 R package (v1.30.1, RRID:SCR_015687), and differential expression analysis was performed using DESeq2. Genes with less than 5 reads in total across all samples were considered not expressed and removed. Differentially expressed genes were called using Benjamini-Hochberg adjusted p-value ≤0.05 and log_2_FC ≥0.5 or ≤−0.5. Over representation analysis (ORA) was performed using clusterProfiler (v4.2.2, RRID:SCR_016884), on the Encode and ChEA consensus transcription factor gene set from the Enrichr web server. Heatmaps depicting ORA analysis were generated using the pheatmap (v1.0.12, RRID:SCR_016418) R package.

## REFERENCES

1. Hargreaves, D.C., and Crabtree, G.R. (2011). ATP-dependent chromatin remodeling: genetics, genomics and mechanisms. Cell Res 21, 396–420. 10.1038/cr.2011.32.

2. Eustermann, S., Patel, A.B., Hopfner, K.P., He, Y., and Korber, P. (2024). Energy-driven genome regulation by ATP-dependent chromatin remodellers. Nat Rev Mol Cell Biol 25, 309–332. 10.1038/s41580-023-00683-y.

3. He, S., Wu, Z., Tian, Y., Yu, Z., Yu, J., Wang, X., Li, J., Liu, B., and Xu, Y. (2020). Structure of nucleosome-bound human BAF complex. Science 367, 875–881. 10.1126/science.aaz9761.

4. Mashtalir, N., Suzuki, H., Farrell, D.P., Sankar, A., Luo, J., Filipovski, M., D’Avino, A.R., St Pierre, R., Valencia, A.M., Onikubo, T., et al. (2020). A Structural Model of the Endogenous Human BAF Complex Informs Disease Mechanisms. Cell 183, 802–817 e824. 10.1016/j.cell.2020.09.051.

5. Yuan, J., Chen, K., Zhang, W., and Chen, Z. (2022). Structure of human chromatin-remodelling PBAF complex bound to a nucleosome. Nature 605, 166–171. 10.1038/s41586-022-04658-5.

6. Mashtalir, N., D’Avino, A.R., Michel, B.C., Luo, J., Pan, J., Otto, J.E., Zullow, H.J., McKenzie, Z.M., Kubiak, R.L., St Pierre, R., et al. (2018). Modular Organization and Assembly of SWI/SNF Family Chromatin Remodeling Complexes. Cell 175, 1272–1288 e1220. 10.1016/j.cell.2018.09.032.

7. Mashtalir, N., Dao, H.T., Sankar, A., Liu, H., Corin, A.J., Bagert, J.D., Ge, E.J., D’Avino, A.R., Filipovski, M., Michel, B.C., et al. (2021). Chromatin landscape signals differentially dictate the activities of mSWI/SNF family complexes. Science 373, 306–315. 10.1126/science.abf8705.

8. Ahmad, K., Brahma, S., and Henikoff, S. (2024). Epigenetic pioneering by SWI/SNF family remodelers. Mol Cell 84, 194–201. 10.1016/j.molcel.2023.10.045.

9. Brahma, S., and Henikoff, S. (2024). The BAF chromatin remodeler synergizes with RNA polymerase II and transcription factors to evict nucleosomes. Nat Genet 56, 100–111. 10.1038/s41588-023-01603-8.

10. Barisic, D., Stadler, M.B., Iurlaro, M., and Schubeler, D. (2019). Mammalian ISWI and SWI/SNF selectively mediate binding of distinct transcription factors. Nature 569, 136–140. 10.1038/s41586-019-1115-5.

11. Schick, S., Grosche, S., Kohl, K.E., Drpic, D., Jaeger, M.G., Marella, N.C., Imrichova, H., Lin, J.G., Hofstatter, G., Schuster, M., et al. (2021). Acute BAF perturbation causes immediate changes in chromatin accessibility. Nat Genet 53, 269–278. 10.1038/s41588-021-00777-3.

12. Iurlaro, M., Stadler, M.B., Masoni, F., Jagani, Z., Galli, G.G., and Schubeler, D. (2021). Mammalian SWI/SNF continuously restores local accessibility to chromatin. Nat Genet 53, 279–287. 10.1038/s41588-020-00768-w.

13. Basurto-Cayuela, L., Guerrero-Martinez, J.A., Gomez-Marin, E., Sanchez-Escabias, E., Escano-Maestre, M., Ceballos-Chavez, M., and Reyes, J.C. (2024). SWI/SNF-dependent genes are defined by their chromatin landscape. Cell Rep 43, 113855. 10.1016/j.celrep.2024.113855.

14. Martin, B.J.E., Ablondi, E.F., Goglia, C., Mimoso, C.A., Espinel-Cabrera, P.R., and Adelman, K. (2023). Global identification of SWI/SNF targets reveals compensation by EP400. Cell 186, 5290–5307 e5226. 10.1016/j.cell.2023.10.006.

15. Kadoch, C., Williams, R.T., Calarco, J.P., Miller, E.L., Weber, C.M., Braun, S.M., Pulice, J.L., Chory, E.J., and Crabtree, G.R. (2017). Dynamics of BAF-Polycomb complex opposition on heterochromatin in normal and oncogenic states. Nat Genet 49, 213–222. 10.1038/ng.3734.

16. Weber, C.M., Hafner, A., Kirkland, J.G., Braun, S.M.G., Stanton, B.Z., Boettiger, A.N., and Crabtree, G.R. (2021). mSWI/SNF promotes Polycomb repression both directly and through genome-wide redistribution. Nat Struct Mol Biol 28, 501–511. 10.1038/s41594-021-00604-7.

17. Nakayama, R.T., Pulice, J.L., Valencia, A.M., McBride, M.J., McKenzie, Z.M., Gillespie, M.A., Ku, W.L., Teng, M., Cui, K., Williams, R.T., et al. (2017). SMARCB1 is required for widespread BAF complex-mediated activation of enhancers and bivalent promoters. Nat Genet 49, 1613–1623. 10.1038/ng.3958.

18. Kadoch, C., Hargreaves, D.C., Hodges, C., Elias, L., Ho, L., Ranish, J., and Crabtree, G.R. (2013). Proteomic and bioinformatic analysis of mammalian SWI/SNF complexes identifies extensive roles in human malignancy. Nat Genet 45, 592–601. 10.1038/ng.2628.

19. Shain, A.H., and Pollack, J.R. (2013). The spectrum of SWI/SNF mutations, ubiquitous in human cancers. PLoS One 8, e55119. 10.1371/journal.pone.0055119.

20. Mittal, P., and Roberts, C.W.M. (2020). The SWI/SNF complex in cancer - biology, biomarkers and therapy. Nat Rev Clin Oncol 17, 435–448. 10.1038/s41571-020-0357-3.

21. Hodis, E., Watson, I.R., Kryukov, G.V., Arold, S.T., Imielinski, M., Theurillat, J.P., Nickerson, E., Auclair, D., Li, L., Place, C., et al. (2012). A landscape of driver mutations in melanoma. Cell 150, 251–263. 10.1016/j.cell.2012.06.024.

22. Shain, A.H., Joseph, N.M., Yu, R., Benhamida, J., Liu, S., Prow, T., Ruben, B., North, J., Pincus, L., Yeh, I., et al. (2018). Genomic and Transcriptomic Analysis Reveals Incremental Disruption of Key Signaling Pathways during Melanoma Evolution. Cancer Cell 34, 45–55 e44. 10.1016/j.ccell.2018.06.005.

23. Tang, J., Fewings, E., Chang, D., Zeng, H., Liu, S., Jorapur, A., Belote, R.L., McNeal, A.S., Tan, T.M., Yeh, I., et al. (2020). The genomic landscapes of individual melanocytes from human skin. Nature 586, 600–605. 10.1038/s41586-020-2785-8.

24. Martinez-Jimenez, F., Muinos, F., Sentis, I., Deu-Pons, J., Reyes-Salazar, I., Arnedo-Pac, C., Mularoni, L., Pich, O., Bonet, J., Kranas, H., et al. (2020). A compendium of mutational cancer driver genes. Nat Rev Cancer 20, 555–572. 10.1038/s41568-020-0290-x.

25. Cerami, E., Gao, J., Dogrusoz, U., Gross, B.E., Sumer, S.O., Aksoy, B.A., Jacobsen, A., Byrne, C.J., Heuer, M.L., Larsson, E., et al. (2012). The cBio cancer genomics portal: an open platform for exploring multidimensional cancer genomics data. Cancer Discov 2, 401–404. 10.1158/2159-8290.CD-12-0095.

26. Gao, J., Aksoy, B.A., Dogrusoz, U., Dresdner, G., Gross, B., Sumer, S.O., Sun, Y., Jacobsen, A., Sinha, R., Larsson, E., et al. (2013). Integrative analysis of complex cancer genomics and clinical profiles using the cBioPortal. Sci Signal 6, pl1. 10.1126/scisignal.2004088.

27. Cancer Genome Atlas, N. (2015). Genomic Classification of Cutaneous Melanoma. Cell 161, 1681–1696. 10.1016/j.cell.2015.05.044.

28. In, G.K., Poorman, K., Saul, M., O’Day, S., Farma, J.M., Olszanski, A.J., Gordon, M.S., Thomas, J.S., Eisenberg, B., Flaherty, L., et al. (2020). Molecular profiling of melanoma brain metastases compared to primary cutaneous melanoma and to extracranial metastases. Oncotarget 11, 3118–3128. 10.18632/oncotarget.27686.

29. Varaljai, R., Horn, S., Sucker, A., Piercianek, D., Schmitt, V., Carpinteiro, A., Becker, K.A., Reifenberger, J., Roesch, A., Felsberg, J., et al. (2021). Integrative Genomic Analyses of Patient-Matched Intracranial and Extracranial Metastases Reveal a Novel Brain-Specific Landscape of Genetic Variants in Driver Genes of Malignant Melanoma. Cancers (Basel) 13. 10.3390/cancers13040731.

30. Carcamo, S., Nguyen, C.B., Grossi, E., Filipescu, D., Alpsoy, A., Dhiman, A., Sun, D., Narang, S., Imig, J., Martin, T.C., et al. (2022). Altered BAF occupancy and transcription factor dynamics in PBAF-deficient melanoma. Cell Rep 39, 110637. 10.1016/j.celrep.2022.110637.

31. Ang, Y.S., Tsai, S.Y., Lee, D.F., Monk, J., Su, J., Ratnakumar, K., Ding, J., Ge, Y., Darr, H., Chang, B., et al. (2011). Wdr5 mediates self-renewal and reprogramming via the embryonic stem cell core transcriptional network. Cell 145, 183–197. 10.1016/j.cell.2011.03.003.

32. Gu, B., and Lee, M.G. (2013). Histone H3 lysine 4 methyltransferases and demethylases in self-renewal and differentiation of stem cells. Cell Biosci 3, 39. 10.1186/2045-3701-3-39.

33. Ernst, J., and Kellis, M. (2017). Chromatin-state discovery and genome annotation with ChromHMM. Nat Protoc 12, 2478–2492. 10.1038/nprot.2017.124.

34. Gaspar-Maia, A., Alajem, A., Meshorer, E., and Ramalho-Santos, M. (2011). Open chromatin in pluripotency and reprogramming. Nat Rev Mol Cell Biol 12, 36–47. 10.1038/nrm3036.

35. Lessard, J., Wu, J.I., Ranish, J.A., Wan, M., Winslow, M.M., Staahl, B.T., Wu, H., Aebersold, R., Graef, I.A., and Crabtree, G.R. (2007). An essential switch in subunit composition of a chromatin remodeling complex during neural development. Neuron 55, 201–215. 10.1016/j.neuron.2007.06.019.

36. Billaud, M., and Santoro, M. (2011). Is Co-option a prevailing mechanism during cancer progression? Cancer Res 71, 6572–6575. 10.1158/0008-5472.CAN-11-2158.

37. Logotheti, S., Marquardt, S., Richter, C., Sophie Hain, R., Murr, N., Takan, I., Pavlopoulou, A., and Putzer, B.M. (2020). Neural Networks Recapitulation by Cancer Cells Promotes Disease Progression: A Novel Role of p73 Isoforms in Cancer-Neuronal Crosstalk. Cancers (Basel) 12. 10.3390/cancers12123789.

38. Venkatesh, H.S., Morishita, W., Geraghty, A.C., Silverbush, D., Gillespie, S.M., Arzt, M., Tam, L.T., Espenel, C., Ponnuswami, A., Ni, L., et al. (2019). Electrical and synaptic integration of glioma into neural circuits. Nature 573, 539–545. 10.1038/s41586-019-1563-y.

39. Biermann, J., Melms, J.C., Amin, A.D., Wang, Y., Caprio, L.A., Karz, A., Tagore, S., Barrera, I., Ibarra-Arellano, M.A., Andreatta, M., et al. (2022). Dissecting the treatment-naive ecosystem of human melanoma brain metastasis. Cell 185, 2591–2608 e2530. 10.1016/j.cell.2022.06.007.

40. Papillon, J.P.N., Nakajima, K., Adair, C.D., Hempel, J., Jouk, A.O., Karki, R.G., Mathieu, S., Mobitz, H., Ntaganda, R., Smith, T., et al. (2018). Discovery of Orally Active Inhibitors of Brahma Homolog (BRM)/SMARCA2 ATPase Activity for the Treatment of Brahma Related Gene 1 (BRG1)/SMARCA4-Mutant Cancers. J Med Chem 61, 10155–10172. 10.1021/acs.jmedchem.8b01318.

41. Chong, J.A., Tapia-Ramirez, J., Kim, S., Toledo-Aral, J.J., Zheng, Y., Boutros, M.C., Altshuller, Y.M., Frohman, M.A., Kraner, S.D., and Mandel, G. (1995). REST: a mammalian silencer protein that restricts sodium channel gene expression to neurons. Cell 80, 949–957. 10.1016/0092-8674(95)90298-8.

42. Hwang, J.Y., and Zukin, R.S. (2018). REST, a master transcriptional regulator in neurodegenerative disease. Curr Opin Neurobiol 48, 193–200. 10.1016/j.conb.2017.12.008.

43. Ooi, L., and Wood, I.C. (2007). Chromatin crosstalk in development and disease: lessons from REST. Nat Rev Genet 8, 544–554. 10.1038/nrg2100.

44. Schoenherr, C.J., and Anderson, D.J. (1995). The neuron-restrictive silencer factor (NRSF): a coordinate repressor of multiple neuron-specific genes. Science 267, 1360–1363. 10.1126/science.7871435.

45. Sehgal, P., and Chaturvedi, P. (2023). Chromatin and Cancer: Implications of Disrupted Chromatin Organization in Tumorigenesis and Its Diversification. Cancers (Basel) 15. 10.3390/cancers15020466.

46. Bergwell, M., Park, J., and Kirkland, J.G. (2024). Differential Modulation of Polycomb-Associated Histone Marks by cBAF, pBAF, and gBAF Complexes. bioRxiv. 10.1101/2023.09.23.557848.

47. Wang, L., Yu, J., Yu, Z., Wang, Q., Li, W., Ren, Y., Chen, Z., He, S., and Xu, Y. (2022). Structure of nucleosome-bound human PBAF complex. Nat Commun 13, 7644. 10.1038/s41467-022-34859-5.

48. Cheli, Y., Ohanna, M., Ballotti, R., and Bertolotto, C. (2010). Fifteen-year quest for microphthalmia-associated transcription factor target genes. Pigment Cell Melanoma Res 23, 27–40. 10.1111/j.1755-148X.2009.00653.x.

49. Seberg, H.E., Van Otterloo, E., and Cornell, R.A. (2017). Beyond MITF: Multiple transcription factors directly regulate the cellular phenotype in melanocytes and melanoma. Pigment Cell Melanoma Res 30, 454–466. 10.1111/pcmr.12611.

50. Stadler, M.B., Murr, R., Burger, L., Ivanek, R., Lienert, F., Scholer, A., van Nimwegen, E., Wirbelauer, C., Oakeley, E.J., Gaidatzis, D., et al. (2011). DNA-binding factors shape the mouse methylome at distal regulatory regions. Nature 480, 490–495. 10.1038/nature10716.

51. Iurlaro, M., Masoni, F., Flyamer, I.M., Wirbelauer, C., Iskar, M., Burger, L., Giorgetti, L., and Schubeler, D. (2024). Systematic assessment of ISWI subunits shows that NURF creates local accessibility for CTCF. Nat Genet. 10.1038/s41588-024-01767-x.

52. Jin, L., Liu, Y., Wu, Y., Huang, Y., and Zhang, D. (2023). REST Is Not Resting: REST/NRSF in Health and Disease. Biomolecules 13. 10.3390/biom13101477.

53. Erickson, C.A. (1993). From the crest to the periphery: control of pigment cell migration and lineage segregation. Pigment Cell Res 6, 336–347. 10.1111/j.1600-0749.1993.tb00611.x.

54. Aoki, H., Hara, A., and Kunisada, T. (2015). White spotting phenotype induced by targeted REST disruption during neural crest specification to a melanocyte cell lineage. Genes Cells 20, 439–449. 10.1111/gtc.12235.

55. Bajpai, R., Chen, D.A., Rada-Iglesias, A., Zhang, J., Xiong, Y., Helms, J., Chang, C.P., Zhao, Y., Swigut, T., and Wysocka, J. (2010). CHD7 cooperates with PBAF to control multipotent neural crest formation. Nature 463, 958–962. 10.1038/nature08733.

56. Torroglosa, A., Villalba-Benito, L., Luzon-Toro, B., Fernandez, R.M., Antinolo, G., and Borrego, S. (2019). Epigenetic Mechanisms in Hirschsprung Disease. Int J Mol Sci 20. 10.3390/ijms20133123.

57. Tagore, M., Hergenreder, E., Perlee, S.C., Cruz, N.M., Menocal, L., Suresh, S., Chan, E., Baron, M., Melendez, S., Dave, A., et al. (2023). GABA Regulates Electrical Activity and Tumor Initiation in Melanoma. Cancer Discov 13, 2270–2291. 10.1158/2159-8290.CD-23-0389.

58. Wang, H., Zheng, Q., Lu, Z., Wang, L., Ding, L., Xia, L., Zhang, H., Wang, M., Chen, Y., and Li, G. (2021). Role of the nervous system in cancers: a review. Cell Death Discov 7, 76. 10.1038/s41420-021-00450-y.

59. Fontanals-Cirera, B., Hasson, D., Vardabasso, C., Di Micco, R., Agrawal, P., Chowdhury, A., Gantz, M., de Pablos-Aragoneses, A., Morgenstern, A., Wu, P., et al. (2017). Harnessing BET Inhibitor Sensitivity Reveals AMIGO2 as a Melanoma Survival Gene. Mol Cell 68, 731–744 e739. 10.1016/j.molcel.2017.11.004.

60. Vardabasso, C., Gaspar-Maia, A., Hasson, D., Punzeler, S., Valle-Garcia, D., Straub, T., Keilhauer, E.C., Strub, T., Dong, J., Panda, T., et al. (2015). Histone Variant H2A.Z.2 Mediates Proliferation and Drug Sensitivity of Malignant Melanoma. Mol Cell 59, 75–88. 10.1016/j.molcel.2015.05.009.

61. Alpsoy, A., and Dykhuizen, E.C. (2018). Glioma tumor suppressor candidate region gene 1 (GLTSCR1) and its paralog GLTSCR1-like form SWI/SNF chromatin remodeling subcomplexes. J Biol Chem 293, 3892–3903. 10.1074/jbc.RA117.001065.

62. Corces, M.R., Trevino, A.E., Hamilton, E.G., Greenside, P.G., Sinnott-Armstrong, N.A., Vesuna, S., Satpathy, A.T., Rubin, A.J., Montine, K.S., Wu, B., et al. (2017). An improved ATAC-seq protocol reduces background and enables interrogation of frozen tissues. Nat Methods 14, 959–962. 10.1038/nmeth.4396.

63. Bolger, A.M., Lohse, M., and Usadel, B. (2014). Trimmomatic: a flexible trimmer for Illumina sequence data. Bioinformatics 30, 2114–2120. 10.1093/bioinformatics/btu170.

64. Gaspar, J.M. (2018). NGmerge: merging paired-end reads via novel empirically-derived models of sequencing errors. BMC Bioinformatics 19, 536. 10.1186/s12859-018-2579-2.

65. Langmead, B., and Salzberg, S.L. (2012). Fast gapped-read alignment with Bowtie 2. Nat Methods 9, 357–359. 10.1038/nmeth.1923.

66. Danecek, P., Bonfield, J.K., Liddle, J., Marshall, J., Ohan, V., Pollard, M.O., Whitwham, A., Keane, T., McCarthy, S.A., Davies, R.M., and Li, H. (2021). Twelve years of SAMtools and BCFtools. Gigascience 10. 10.1093/gigascience/giab008.

67. Zhang, Y., Liu, T., Meyer, C.A., Eeckhoute, J., Johnson, D.S., Bernstein, B.E., Nusbaum, C., Myers, R.M., Brown, M., Li, W., and Liu, X.S. (2008). Model-based analysis of ChIP-Seq (MACS). Genome Biol 9, R137. 10.1186/gb-2008-9-9-r137.

68. Zang, C., Schones, D.E., Zeng, C., Cui, K., Zhao, K., and Peng, W. (2009). A clustering approach for identification of enriched domains from histone modification ChIP-Seq data. Bioinformatics 25, 1952–1958. 10.1093/bioinformatics/btp340.

69. Ramirez, F., Ryan, D.P., Gruning, B., Bhardwaj, V., Kilpert, F., Richter, A.S., Heyne, S., Dundar, F., and Manke, T. (2016). deepTools2: a next generation web server for deep-sequencing data analysis. Nucleic Acids Res 44, W160–165. 10.1093/nar/gkw257.

70. Quinlan, A.R., and Hall, I.M. (2010). BEDTools: a flexible suite of utilities for comparing genomic features. Bioinformatics 26, 841–842. 10.1093/bioinformatics/btq033.

71. Wang, Q., Li, M., Wu, T., Zhan, L., Li, L., Chen, M., Xie, W., Xie, Z., Hu, E., Xu, S., and Yu, G. (2022). Exploring Epigenomic Datasets by ChIPseeker. Curr Protoc 2, e585. 10.1002/cpz1.585.

72. Ernst, J., and Kellis, M. (2012). ChromHMM: automating chromatin-state discovery and characterization. Nat Methods 9, 215–216. 10.1038/nmeth.1906.

73. Ross-Innes, C.S., Stark, R., Teschendorff, A.E., Holmes, K.A., Ali, H.R., Dunning, M.J., Brown, G.D., Gojis, O., Ellis, I.O., Green, A.R., et al. (2012). Differential oestrogen receptor binding is associated with clinical outcome in breast cancer. Nature 481, 389–393. 10.1038/nature10730.

74. Dobin, A., Davis, C.A., Schlesinger, F., Drenkow, J., Zaleski, C., Jha, S., Batut, P., Chaisson, M., and Gingeras, T.R. (2013). STAR: ultrafast universal RNA-seq aligner. Bioinformatics 29, 15–21. 10.1093/bioinformatics/bts635.

75. Liao, Y., Smyth, G.K., and Shi, W. (2019). The R package Rsubread is easier, faster, cheaper and better for alignment and quantification of RNA sequencing reads. Nucleic Acids Res 47, e47. 10.1093/nar/gkz114.

76. Love, M.I., Huber, W., and Anders, S. (2014). Moderated estimation of fold change and dispersion for RNA-seq data with DESeq2. Genome Biol 15, 550. 10.1186/s13059-014-0550-8.

77. Kuleshov, M.V., Jones, M.R., Rouillard, A.D., Fernandez, N.F., Duan, Q., Wang, Z., Koplev, S., Jenkins, S.L., Jagodnik, K.M., Lachmann, A., et al. (2016). Enrichr: a comprehensive gene set enrichment analysis web server 2016 update. Nucleic Acids Res 44, W90–97. 10.1093/nar/gkw377.

78. Mounir, M., Lucchetta, M., Silva, T.C., Olsen, C., Bontempi, G., Chen, X., Noushmehr, H., Colaprico, A., and Papaleo, E. (2019). New functionalities in the TCGAbiolinks package for the study and integration of cancer data from GDC and GTEx. PLoS Comput Biol 15, e1006701. 10.1371/journal.pcbi.1006701.

