## Supplemental Figures 1-6 for "The SWI/SNF PBAF complex facilitates REST occupancy at repressive chromatin"

Grossi *et al.*, Fig. S1

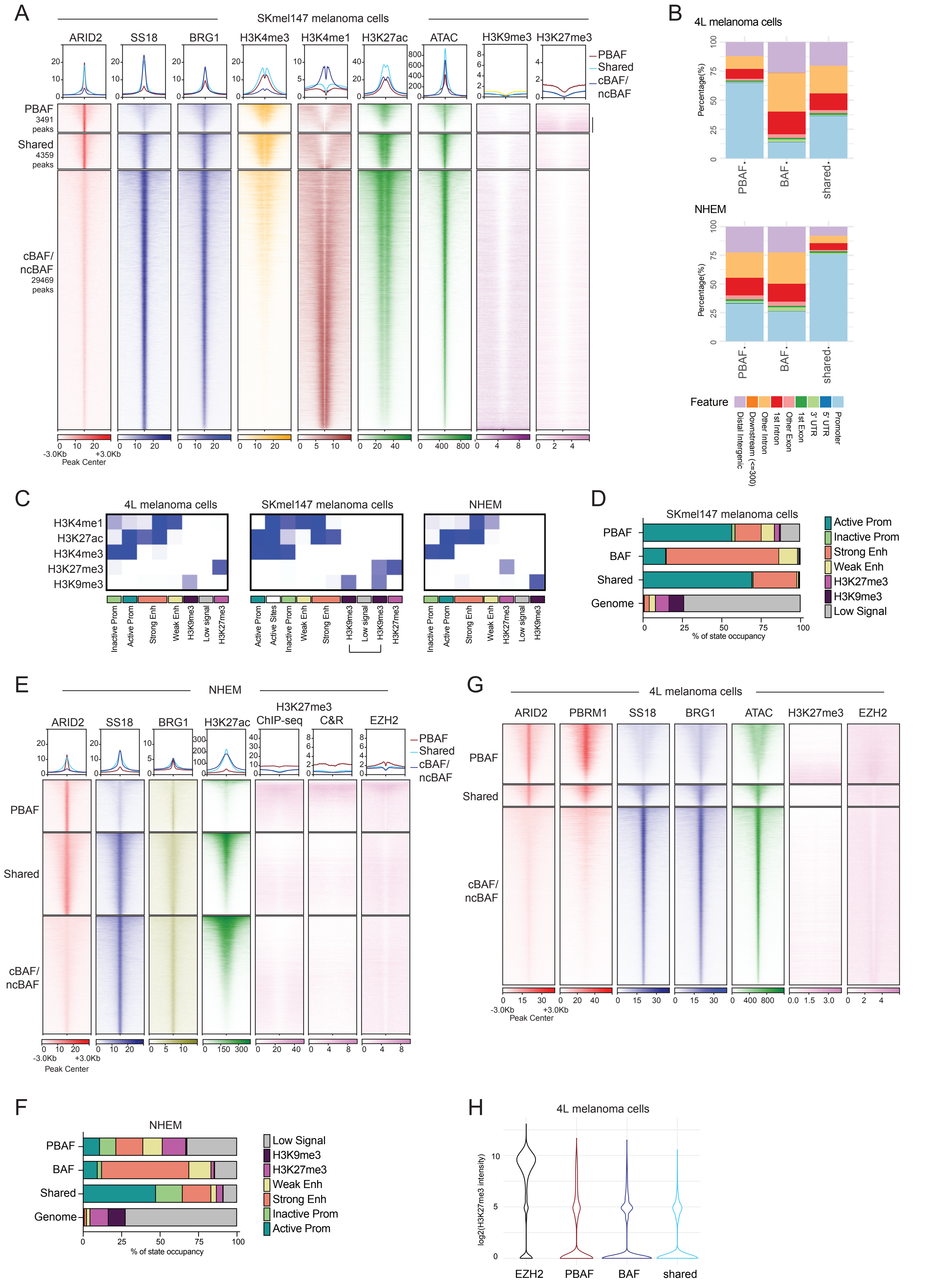

**Figure S1. A subset of PBAF-enriched regions associates with repressed chromatin.** (A) Heatmap displaying the genomic distribution of SWI/SNF components and histone marks, at PBAF-, Shared- or BAF- (cBAF/ncBAF-) enriched regions sorted by H3K27ac signal in SKmel147 melanoma cells. Black sidebar highlights H3K27me3-rich PBAF regions. (B) Bar plots depicting the percentage of PBAF-, shared and BAF-enriched regions occupying proximal or distal regulatory regions in 4L cells or NHEM. (C) ChromHMM emission model generated using epigenomic data of histone marks in 4L, SKmel147 and NHEM cells. Clusters presenting H3K9me3-only and H3K27me3/H3K9me3 signal in SKmel147 were grouped for consistency with 4L and NHEM data (see Methods). (D) Distribution of chromatin states associated with different subcomplexes and compared to their distribution in the genome in SKmel147 cells. (E) Heatmap comparing H3K27me3 signal obtained using ChIP-seq or CUT&RUN at PBAF-, shared and BAF-enriched regions in NHEM. EZH2 heatmap relative to metagene plot in Fig. 1D (F) Distribution of chromatin states associated with different SWI/SNF subcomplexes in NHEM as in Fig. 1C, using H3K27me3 ChIP-seq data. (G) Heatmaps relative to metagene plot in Fig. 1D (H) Violin plot depicting H3K27me3 distribution in 4L cells at all EZH2-, PBAF-, BAF- enriched or shared regions, regardless of their overlap with H3K27me3. A pseudocount was added for logarithmic scale visualization.

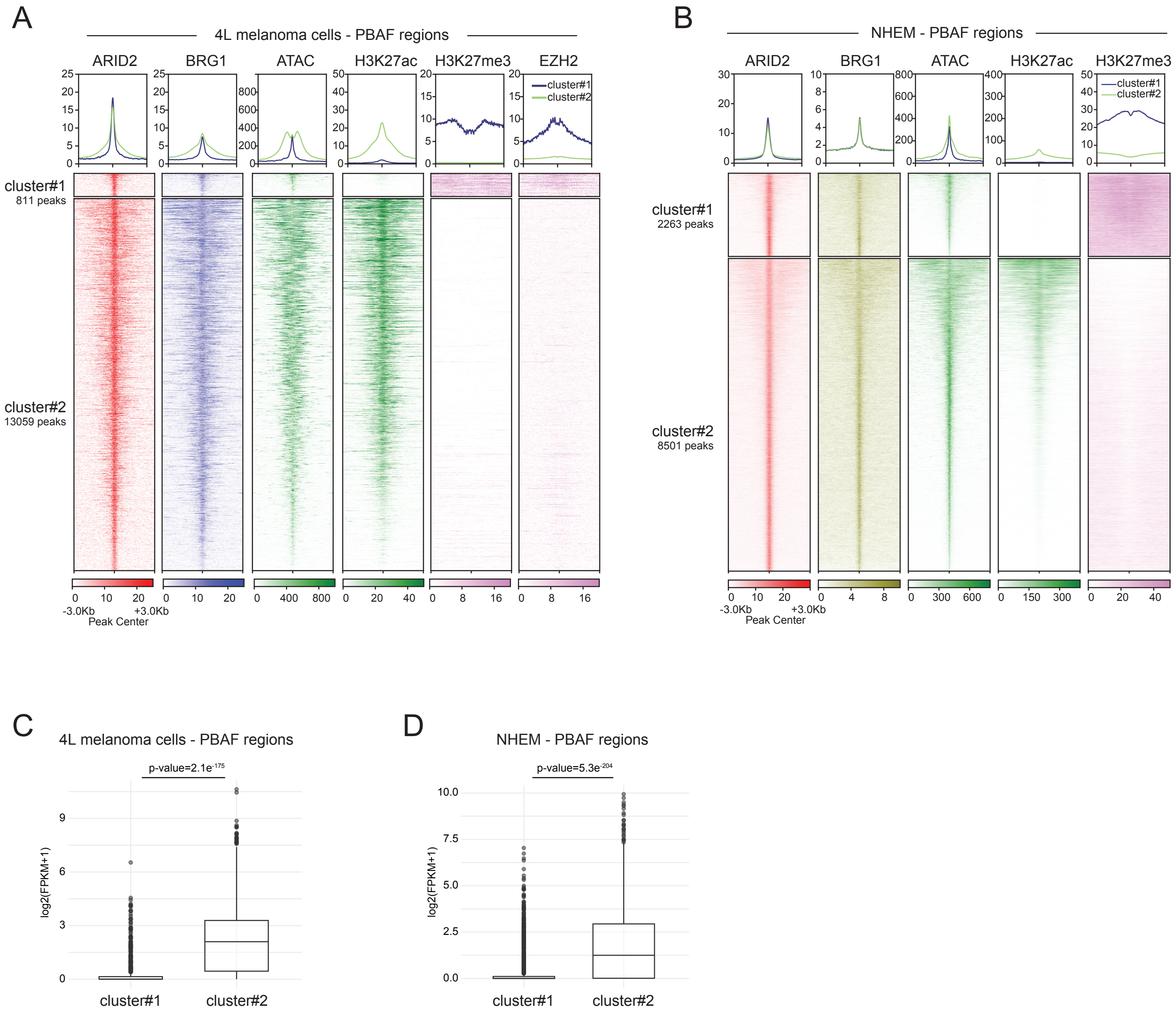


**Figure S2. PBAF-enriched regions present lower chromatin accessibility compared to BAF-enriched sites.** (A-B) Heatmaps displaying PBAF-enriched regions grouped as repressed (cluster#1) or active (cluster #2) sites by k-means clustering in 4L melanoma cells (A) and NHEM (B). (C-D) Boxplot depicting RNA expression levels (FPKM) of genes associated with repressed (cluster#1) or active (cluster#2) PBAF regions, as defined in A-B. In 4L melanoma cells (C) genes in cluster#1 = 633; genes in cluster#2 = 8233. In NHEM (D) genes in cluster#1 = 1635; genes in cluster#2 = 5436. A pseudocount was added for logarithmic scale visualization. Significance was calculated by Mann-Whitney U test.

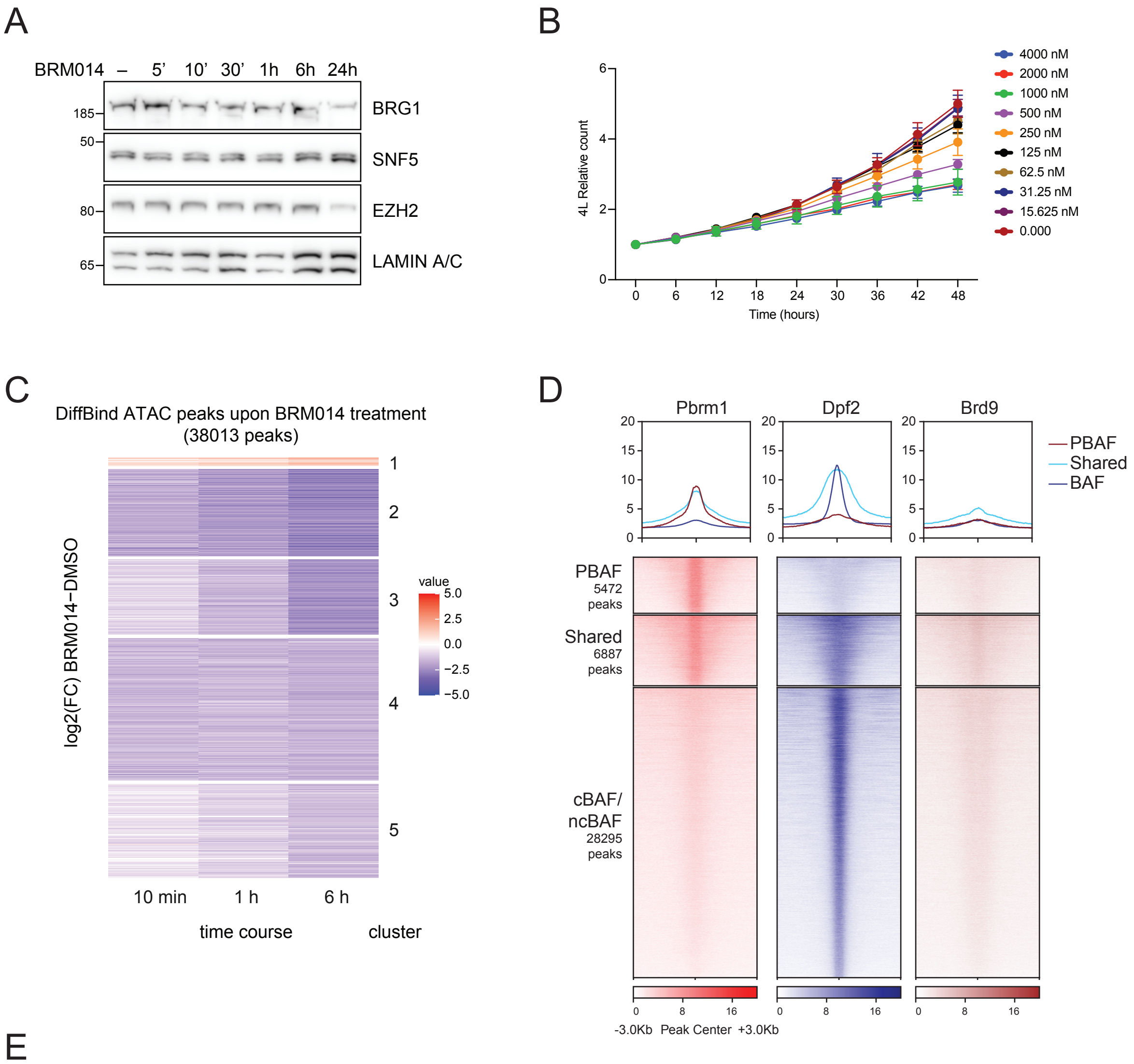


E

| SWI/SNF subcomplex | Motif | Name | p-value | % targets with motif |
|---|---|---|---|---|
| PBAF | 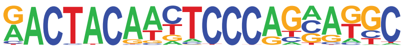 | RONIN/GFY | 1e-92 | 3.62% |
| Shared | 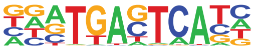 | AP-1 family | 1e-5415 | 36.49% |
| BAF | 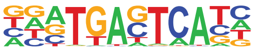 | AP-1 family | 1e-59 | 7.14% |

**Figure S3. SWI/SNF ATPase inhibition affects chromatin accessibility in melanoma cells.** (A) Western blot analysis of 4L chromatin extracts upon 1 µM BRM014 treatment. Control cells were treated with DMSO (–). LAMIN A/C levels were used as loading control. (B) Growth analysis of 4L melanoma cells subject to different concentrations of BRM014 compound over a 48-hour time course. Cell counts for each time point were normalized to the corresponding time 0. A concentration of 1 µM was used for subsequent ATAC-seq studies. (C) Heatmap displaying significant changes (FDR<0.05; $|\log_2 FC>1|$) upon treatment with 1 µM BRM014 at different time points compared to DMSO-treated control 4L melanoma cells. ATAC sites are clustered into five groups by k-means clustering. (D) Heatmap displaying the genomic distribution of SWI/SNF components , at PBAF-, Shared- or BAF- (cBAF/ncBAF-) enriched regions in murine NMuMG epithelial cells[13]. (E) Motif enrichment analysis of PBAF-, shared or BAF, enriched regions (as defined in D) in NMuMG cells. Only top-enriched TF families are shown.

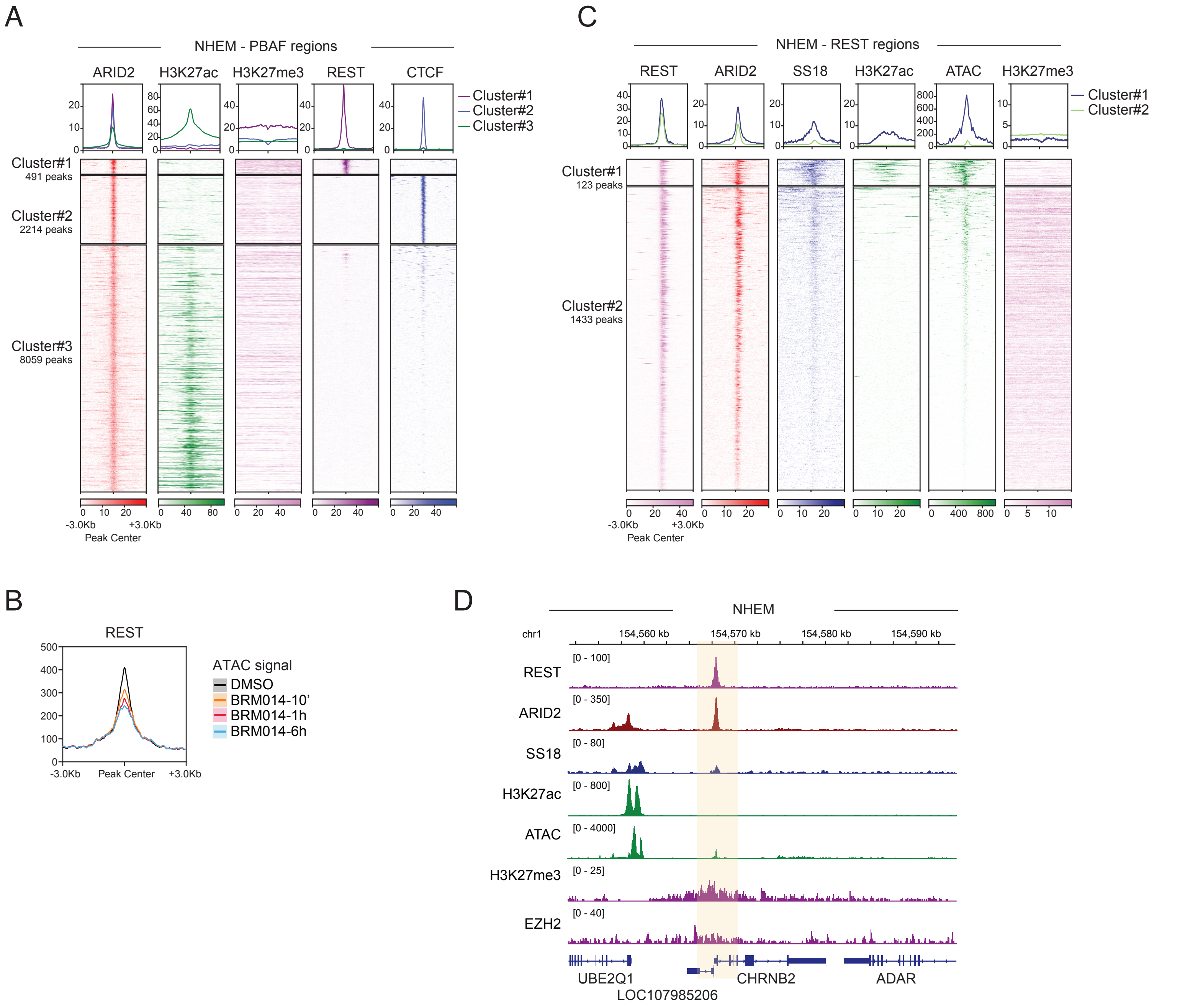


**Figure S4. REST is preferentially associated to the PBAF complex.** (A) Heatmap displaying different TF occupancy at PBAF-enriched regions in NHEM. (B) Heatmap showing distribution of SWI/SNF components and histone marks at REST regions in NHEM. REST sites were grouped into active (cluster#1) or repressed (cluster #2) sites by k-means clustering. (C) Metagene profile illustrating changes in chromatin accessibility at REST regions, as measured by ATAC-seq at different time points of 1 μM BRM014 treatment in 4L melanoma cells. (D) Genomic browser snapshot illustrating occupancy of SWI/SNF and PRC2 components, histone marks and REST at *CHRNB2* neuronal gene in NHEM.

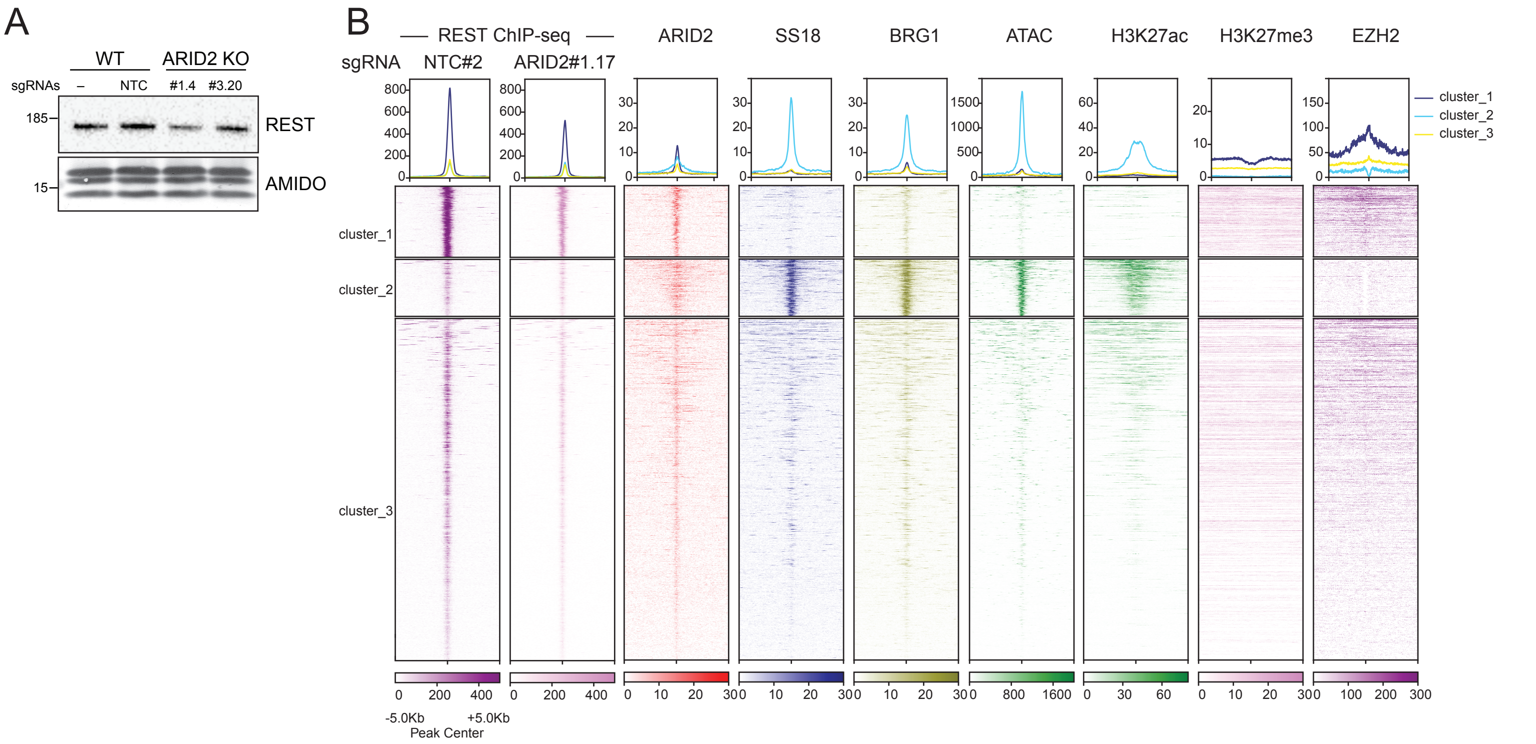


**Figure S5. PBAF depletion impairs REST binding at repressed regions.** (A) Western blot analysis of SKmel147 chromatin extracts upon ARID2 depletion using two different sgRNAs. NTC, non-targeting control. Amido black is used as loading control (B) Heatmap displaying changes in REST binding upon ARID2 KO in 4L melanoma cells. K-means clustering was employed to group different regions based on their chromatin features. Genomic occupancy of SWI/SNF components and histone marks, as well as chromatin accessibility levels is also shown.

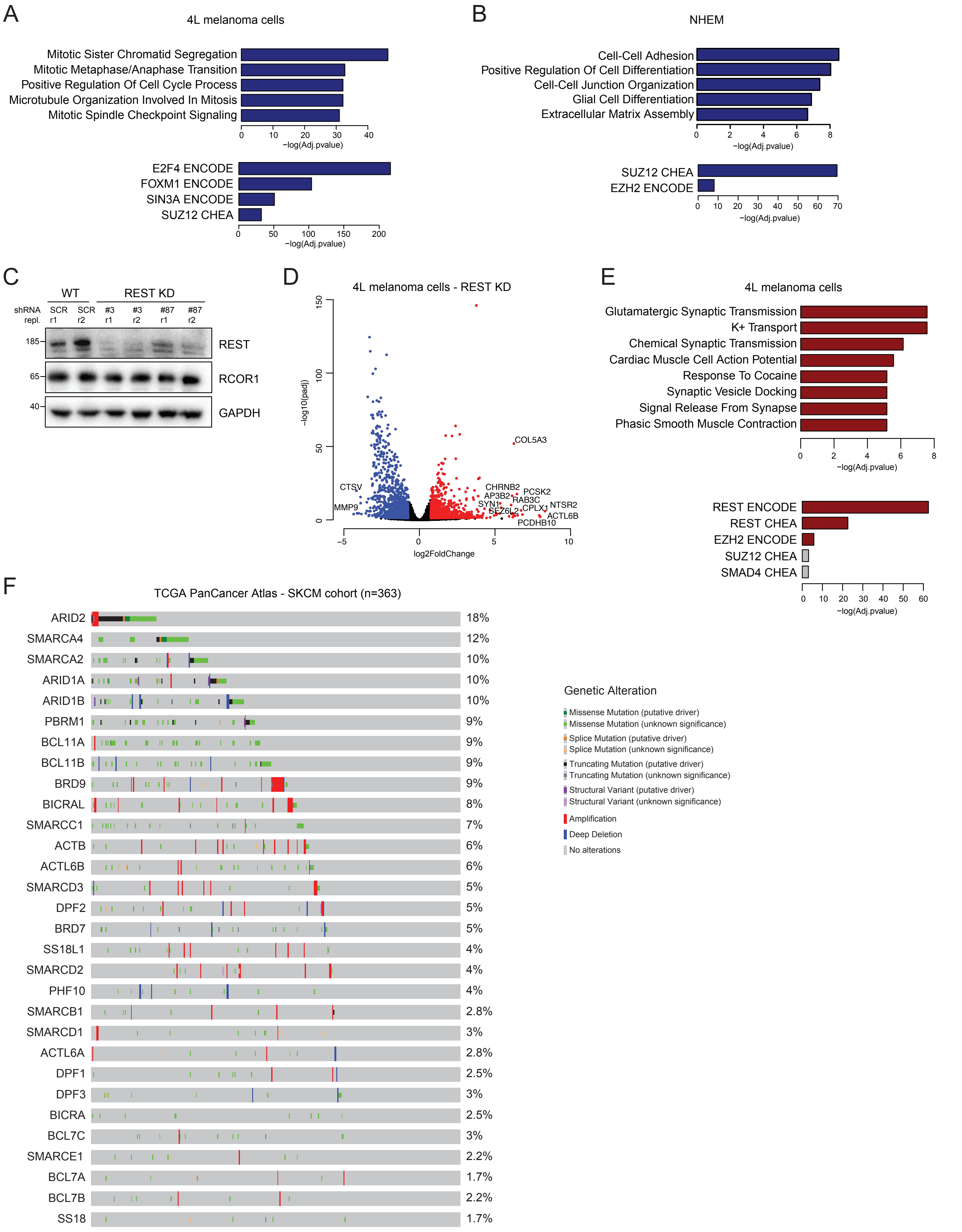


**Figure S6. PBAF loss derepresses REST targets in melanoma cells and patients.** (A-B) Biological processes (top) and TF enrichment (bottom) GO analysis of downregulated genes upon ARID2 KO in 4L cells (A) and NHEM (B). (C) Western blot analysis of whole cell extracts of 4L cells upon REST KD using two independent shRNAs in biological duplicate. REST knockdown does not affect RCOR1 (REST Corepressor 1) levels. GAPDH levels were detected as loading control. (D) Volcano plot displaying gene expression changes upon REST knockdown in 4L melanoma cells (adj.pvalue<0.05; $|log_2FC>0.75|$) (E) Biological processes and TF enrichment GO analysis of common upregulated genes in ARID2 KO and REST KD 4L cells. (F) Panel of genetic alterations found in SWI/SNF components in SKCM patient samples.
